# Widespread mono- and oligoadenylation direct small noncoding RNA maturation versus degradation fates

**DOI:** 10.1101/2025.01.31.635978

**Authors:** Cody Ocheltree, Blake Skrable, Anastasia Pimentel, Timothy Nicholson-Shaw, Suzanne R. Lee, Jens Lykke-Andersen

**Affiliations:** Department of Molecular Biology, School of Biological Sciences, University of California San Diego, La Jolla, CA, 92093, USA

## Abstract

Small non-coding RNAs (sncRNAs) are subject to 3’ end trimming and tailing activities that impact maturation versus degradation decisions during biogenesis. To investigate the dynamics of human sncRNA 3’ end processing at a global level we performed genome-wide 3’ end sequencing of nascently-transcribed and steady-state sncRNAs. This revealed widespread post-transcriptional adenylation of nascent sncRNAs, which came in two distinct varieties. One is characterized by oligoadenylation, which is transient, promoted by TENT4A/4B polymerases, and most commonly observed on unstable snoRNAs that are not fully processed at their 3’ ends. The other is characterized by monoadenylation, which is broadly catalyzed by TENT2 and, in contrast to oligoadenylation, stably accumulates at the 3’-end of sncRNAs, including Polymerase-III-transcribed (Pol-III) RNAs and a subset of small nuclear RNAs. Monoadenylation inhibits Pol-III RNA post-transcriptional 3’ uridine trimming and extension and, in the case of 7SL RNAs, prevents their accumulation with nuclear La protein and promotes their biogenesis towards assembly into cytoplasmic signal recognition particles. Thus, the biogenesis of human sncRNAs involves widespread mono- or oligo-adenylation with divergent impacts on sncRNA fates.

## INTRODUCTION

The vast majority of eukaryotic RNAs undergo post-transcriptional processing critical for their biogenesis into functional mature RNAs. RNA 3’-ends in particular are subject to a wide variety of processing events including exonucleolytic trimming by 3’-to-5’ exonucleases, addition of post-transcriptional nucleotides by polymerases, or 3’ end nucleotide modifications such as 2’,3’ cyclic phosphorylation or 2’O-methylation. Some of the most abundant RNAs in eukaryotic cells are small non-coding RNAs (sncRNAs), which perform a wide variety of critical functions in gene expression. The majority of sncRNAs undergo processing at their 3’ ends after their initial transcription, which can impact sncRNA cellular localization and function. Moreover, 3’ end trimming and tailing towards sncRNA maturation have been observed to occur in competition with degradation from the 3’ end in a process thought to distinguish functional from non-functional molecules (1–5).

SncRNAs can be divided into distinct classes in terms of how their 3’ ends are initially formed. One consists of RNA polymerase II-transcribed sncRNAs, such as spliceosomal snRNAs, whose nascent 3’ ends form via co-transcriptional cleavage by the Integrator complex (6–9). This cleavage results in short encoded 3’-end extensions, which subsequently undergo various degrees of trimming (1, 3, 4). A second class consists of snoRNAs and scaRNAs that are encoded within introns of protein-coding or non-coding genes (10). The 3’ ends of these sncRNAs are formed by the process of splicing, which is followed by intron debranching and trimming to generate the mature molecules (11). A third class consists of RNA Polymerase-III (Pol-III or Pol3)-transcribed RNAs, which are terminated at oligo(T) DNA termination sequences (12–14). The resulting 3’-uridine termini generally undergo subsequent trimming or, in the case of U6 snRNA, extension to generate the mature RNAs (15–18).

A large number of exonucleases and polymerases participate in the processing of sncRNA 3’ ends. Some of these function in RNA maturation while others promote degradation. For example, 3’-to-5’ exonucleases TOE1, PARN, and USB1 promote 3’ end maturation of a large variety of sncRNAs (19), whereas DIS3 exonucleases, either as components of the exosome (DIS3 or DIS3L in human) (20, 21), or acting on its own in the cytoplasm (DIS3L2) (22), promote RNA degradation. 3’ end tailing by polymerases has also been associated with either maturation or degradation dependent on the polymerase and RNA. For example, oligo-uridylation of RNAs by Terminal Uridylyltransferases (TUT) 4 and 7 promote degradation by DIS3L2 in the cytoplasm (23, 24), whereas oligoadenylation of RNAs by Terminal Nucleotidyltransferases TENT4A and TENT4B is associated with degradation by the exosome in the nucleus (25). However, uridylation and adenylation have also been associated with RNA maturation and stability. For example, uridylation in the nucleus by TUT1 promotes maturation of U6 snRNA (26, 27). Moreover, monoadenylation by the Terminal Nucleotidyltransferase TENT2 has been observed to stabilize a subset of microRNAs (28, 29) and oligoadenylation has been proposed to promote 3’ end maturation activities of PARN (2, 30, 31) and TOE1 (1, 3, 4). However, the general principles that govern these 3’ end trimming and tailing events and how they drive the competition between sncRNA maturation and degradation remain poorly defined.

Many Pol-III RNAs undergo trimming or tailing of the initial 3’ oligouridine termini produced during transcription termination. For U6 snRNA, U-tail extension by TUT1 is terminated by a 2’,3’ cyclic phosphorylation event, which is important for maturation and stability (17, 32). Another subset of Pol-III RNAs (as well as some snRNAs) have been observed to undergo monoadenylation to varying degrees (33) and this monoadenylation has been observed to oppose 3’ uridylation in vitro and when injected into Xenopus oocyte nuclei (34). One example is 7SL RNAs, which serve as the central scaffold of the Signal Recognition Particle (SRP) that functions in the co-translational translocation of polypeptides into the endoplasmic reticulum (ER) (35, 36). Biogenesis of 7SL RNAs involves assembly with several SRP proteins in the nucleus prior to nuclear export and association with a final SRP protein component, SRP54, in the cytoplasm (37–39). In mammalian cells, 7SL RNA is known to be processed at the 3’ end with the removal of terminal uridines and the addition of a monoadenosine tail (40, 18, 33), but how this may impact 7SL RNA biogenesis and assembly into the SRP is unknown.

In this study, we investigated the genome-wide dynamics of 3’ end processing of human sncRNAs by sequencing 3’ ends of nascent and steady state sncRNAs, ≈100-500 nucleotides in length. This revealed widespread sncRNA post-transcriptional A-tailing, which was observed in two forms. One consisted of transient oligo(A)-tails, which were observed primarily on partially-processed snoRNAs and scaRNAs, and correlated with instability rather than 3’ end maturation. Another consisted of mono(A)-tailing, which stably accumulated on a majority Pol-III RNA species and a subset of snRNAs. We identified TENT2 as a non-canonical polymerase that broadly monoadenylates Pol-III RNAs and snRNAs and find that mono(A)-tailing by TENT2 inhibits both uridylation and deuridylation of Pol-III RNAs. Moreover, we present evidence that monoadenylation promotes 7SL RNA biogenesis by inhibiting 7SL RNA interaction with nuclear La protein and promoting the assembly into cytoplasmic SRP particles. These findings identify divergent roles of mono- and oligo-adenylation in maturation versus degradation decisions during sncRNA biogenesis.

## MATERIAL AND METHODS

### Mammalian cell culture

All cells were maintained in Dulbecco’s Modified Eagle Medium (DMEM, Gibco) supplemented with 10% Fetal Bovine Serum (FBS, Gibco) and 1% penicillin/streptomycin (Gibco) at 37°C, 5% CO_2_. Mycoplasma testing was routinely performed.

### Global 3’ end sequencing of nascent small RNAs

HEK293 T-REx cells were incubated with either 0.5 mM 5-ethynyl uridine (EU; Thermo Fisher) or an equivalent volume of DMSO for 2 hours in order to accumulate sufficient molecules for detection by global sequencing and harvested in TRIzol reagent (Thermo Fisher). Total RNA was isolated according to the manufacturer’s recommendation. Small RNA from 180 μg of total RNA was isolated by separation in a 9% polyacrylamide/6M urea denaturing gel. After Sybr Gold staining (Thermo Fisher), small RNAs approximately 100 to 500 nucleotides in length were excised and eluted with gel elution buffer (0.3 M sodium acetate pH 5.3, 1 mM EDTA, 0.1% SDS) by end-over-end rotation overnight at 4°C. Eluted small RNAs were purified by RNA clean & concentrator columns (Zymo Research). Genomic DNA was removed using Turbo DNA-free kit (Thermo Fisher) and ribosomal (r)RNAs were depleted using RiboCOP rRNA depletion kit (Lexogen) per manufacturer’s recommendations. RNA samples were treated with FastAP (Thermo Fisher) in 25 μl total volume and subsequently PNK (NEB) in 100 μl total volume in order to remove RNA 5’- and 3’-phosphates. RNA samples were again purified using RNA Clean & Concentrator columns (Zymo Research). AG15N or AG16N RNA adaptors (1 μM, Supplementary Table S1) were ligated to the RNA 3’ ends by incubation at 25°C for 90 minutes in a 40 µl reaction containing 9% DMSO (Sigma), ligase buffer (50 mM Tris-HCl pH 7.5, 10 mM MgCl_2_, 1 mM DTT), 1 mM ATP (Thermo Fisher), 16 units RNaseOUT (Thermo Fisher), 20% PEG 8000 (NEB), and 80 units T4 RNA ligase (NEB). RNA from ligation reactions were purified using RNA Clean & Concentrator columns (Zymo Research). In order to isolate nascent RNAs labeled with EU, a portion of each sample was processed with Click-it nascent RNA capture kit (Thermo Fisher) as previously described (3), while the remaining portion did not undergo nascent enrichment in order to represent the steady state population. To determine the effectiveness of the nascent enrichment step, two truncated exogenous b-globin RNA probes approximately 250 nucleotides in length were spiked in to samples prior to nascent extraction – one probe was *in-vitro* transcribed with a 1:20 ratio of 5-ethynyl-UTP:UTP in order to mimic the estimated nucleotide ratio in cell culture, while a second probe was *in-vitro* transcribed only with UTP (Supplementary Table S1). Reverse transcription of nascently-made RNAs was performed on-bead in a total of 20 µl with 0.5 μM AR17 primer (Supplemental Table 3), 12.5 nM spike-in probe primer (Supplementary Table S1), AffinityScript buffer (1X, Agilent), 10 mM DTT, 4 mM dNTPs, 12 units RNaseOUT (Thermo Fisher), and AffinityScript Reverse Transcriptase (1X, Agilent) at 55°C for 45 minutes, followed by 15 minutes incubation at 70°C and 5 minutes of incubation at 85°C to release cDNA. Reverse transcription of RNA representing steady state accumulation was performed in tandem. Excess primer and RNA were removed from cDNA samples by incubating with 3.5 μl ExoSAP-IT (Thermo Fisher) at 37°C for 15 minutes, then treated with 3 μl of 1M NaOH at 70°C for 12 minutes and subsequently neutralized with 3 μl of 1M HCl. cDNA was extracted with phenol:chloroform:isoamyl alcohol and precipitated with 0.1 volume of 3 M sodium acetate pH 5.3 and 2.5 volumes of ethanol. A 3Tr3 adaptor (Supplementary Table S1) was ligated to cDNA 3’-ends at a final concentration of 3.2 µM in a 20 μl reaction with 5% DMSO (Sigma), in ligase buffer (50 mM Tris-HCl pH 7.5, 10 mM MgCl_2_, 1 mM DTT), 1 mM ATP (Thermo Fisher), and 45 units T4 RNA ligase (NEB) at 25°C for 16 hours. cDNA to be sequenced was amplified in two stages of Polymerase Chain Reaction (PCR) using Q5 DNA polymerase (NEB). For the first PCR reaction, the cDNA library was amplified using 3’ adaptor primer (AR17) and a primer complementary to the 5’ adaptor (RC_3Tr3) (Supplementary Table S1) for 8 cycles. The PCR product was purified by AMPure XP beads (Beckman Coulter) per manufacturer’s recommendation. The second PCR reaction was performed using Illumina Truseq D50X and D70X primers (Supplementary Table S1) for 18 cycles. The library quality was monitored by qPCR for select genes and Tapestation (Agilent) analyses. The relative ratios of EU-labeled to unlabeled spike-in probe cDNA were compared via qPCR in both nascently-extracted samples and samples representing the steady state (Supplementary Table S1). 100 bp paired-end sequencing was performed on an Illumina Novaseq S4.

### Sequencing data analyses

Fastq files were first subjected to 3’ adaptor and PCR duplicate removal using custom python scripts (https://pypi.org/project/jla-demultiplexer/). Residual Illumina adaptor sequence was removed with Cutadapt (41). Reads were mapped to the human genome (version hg38) using STAR 2.7.11b (42). A three-pass alignment strategy against a small RNA genome was used as previously described (4), but with a modification in order to improve the local alignment step which provides post-transcriptional nucleotide modification information. After performing a three-pass end-to-end alignment, which includes a 5’-hard clip of 10 nucleotides to facilitate alignment of reads with post-transcriptional nucleotide modifications, the aligned reads were grouped by gene and re-converted into individual fastq files for each gene detected using bedtools (43) and samtools (44). Individual-gene fastq files were then re-aligned with a three-pass local alignment to single-gene genomic sequences corresponding to the gene each file was previously aligned to in the end-to-end alignment. Gene-specific 3’ end information and graphs were subsequently generated using Tailer (45) (https://github.com/TimNicholsonShaw/tailer) using the global alignment mode. RNAs that were full-length or extended up to 50 nucleotides were evaluated for 3’ end processing and modification, while RNAs truncated >10 base pairs from the mature 3’ end were removed from analyses. Small non-coding RNA species with 1 or more reads in each of three biological replicates in the nascent condition were evaluated in subsequent analyses.

### Gene-specific RNA 3’ end sequencing

RNA was isolated using TRIzol (Thermo Fisher) per manufacturer’s recommendation. RNA was subsequently treated with DNase I (Zymo Research), purified with RNA Clean & Concentrator columns (Zymo Research), and AG15N or AG16N RNA adaptors were ligated to RNA 3’ ends as described above. Following RNA purification with RNA Clean & Concentrator columns (Zymo Research), cDNA was synthesized with Superscript III (Thermo Fisher) using the AR17 primer (Supplementary Table S1). Gene-specific RNA 3’ end sequencing libraries were generated using gene-specific forward primers (Supplementary Table S1) and the AR17 reverse primer (Supplementary Table S1) with Q5 polymerase (NEB). Libraries were sequenced on an Illumina MiSeq platform and analyzed as previously described (3).

### CRISPR/Cas9 knockout cell lines

The human *Terminal Nucleotidyltransferase 2* (*TENT2)* gene was targeted with guide RNAs against PAM sites located in 3’-exon 2 and 5’-intron 2 (Supplementary Figure S4B and Table S4). The guide RNA-containing constructs and Cas9 vector were prepared and transfected as described previously (46) into HEK293 T-REx cells. Cells were selected for transfection construct expression by fluorescence-activated cell sorting for GFP-positive fluorescence. GFP-positive cells were plated as single-cell colonies and evaluated for genomic deletion of *TENT2* sequence using primers flanking exon 2 and intron 2 (Supplementary Table S1). PCR products were cloned into the PX459 plasmid (46), transformed into DH5a *E. coli* and prepared for sanger and nanopore sequencing following plasmid purification (QIAprep, Qiagen). Six sanger sequencing reactions and four nanopore sequencing reactions were performed per knockout clone. Three unique alleles were detected from clone A and two from clone B from ten and six successful sequencing reactions, respectfully (Supplementary Figure S4B and Table S5). The loss of the exon 2 splicing junction is predicted to result in a premature stop codon. A loss of TENT2 protein expression for TENT2 KO clones but not the Control clone was validated by western blotting (Supplementary Figure S4C). A control clone was derived from cells which were GFP-positive, but did not present evidence for TENT2 genomic deletion in alleles from four nanopore sequencing reactions (Supplementary Table S2) or absence of TENT2 protein.

### Western blotting

Western blots were performed by separating proteins in SDS-polyacrylamide gels followed by transfer to nitrocellulose membranes using standard procedures. Membranes were incubated overnight at 4°C with rabbit polyclonal anti-b-tubulin antibody (2146, Cell Signaling Technologies) and rabbit polyclonal anti-TENT2 antibody (PA5-65876, Thermo Fisher) diluted 1:1,000 in TBS with 0.1% Tween 20 (TBST) and 3% bovine serum albumin (BSA). Secondary donkey anti-rabbit antibodies conjugated to HRP (Thermo Fisher) were diluted 1:10,000 in TBST with 3% BSA and incubated for two hours at room temperature. Protein was visualized with SuperSignal West Femto substrate (Thermo Fisher) using an Odyssey Fc imaging system (LI-COR).

### Generation of plasmid constructs and stable cell lines

Gibson assembly (New England Biolabs) was used to insert a cDNA copy of the human *TENT2* gene with a N-terminal-3XFLAG-tag sequence into the pcDNA5/FRT/TO (Thermo Fisher) plasmid, creating pcDNA5-3XFLAG-TENT2WT. The 3XFLAG-tag sequence alone was additionally inserted into pcDNA5/FRT/TO. Site-directed mutagenesis (New England Biolabs) was used to create a silent mutation in the coding sequence of *TENT2*, conferring siRNA resistance and creating pcDNA5-3XFLAG-TENT2WT-R. Site-directed mutagenesis was again used to mutate two codons (ATGGTGA>CTGGTGC), resulting in Asp to Ala mutations at residues 213 and 215 critical for nucleotidyltrasferase catalytic activity of TENT proteins, creating pcDNA5-3XFLAG-TENT2DADA-R (47). pcDNA5-3XFLAG, pcDNA5-3XFLAG-TENT2WT-R and pcDNA5-3XFLAG-TENT2DADA-R constructs were subsequently integrated into the insertion site of Flp-In 293 T-REx cells (Thermo Fisher) per manufacturer’s recommendations in order to generate stable cell lines expressing 3XFLAG, 3XFLAG-TENT2WT-R and 3XFLAG-TENT2DADA-R under control of a tetracycline-inducible promoter. Stable cell lines were treated with 50 ng/ml tetracycline 24 hours prior to harvest in order to induce expression of 3XFLAG, 3XFLAG-TENT2WT-R, and 3XFLAG-TENT2DADA-R proteins. Plasmid sequences are available upon request.

To generate vectors for expression of exogenous 7SL1 RNA, genomic DNA from HeLa cells was isolated with QuickExtract (Biosearch Technologies Inc.) and the 7SL1 gene was amplified by PCR with Q5 polymerase (NEB), 3% DMSO, and primers positioned 154 base pairs upstream of the first transcribed nucleotide and 39 base pairs downstream of the annotated 3’-end (Supplementary Table S1). The amplicon was cloned into the pUC19 plasmid using standard molecular cloning techniques. The 7SL1 sequence was subsequently altered by Q5 (NEB) PCR amplification using primers (Supplementary Table S1) to achieve site-directed mutagenesis in order to create a distinguishable nucleotide barcode from endogenous 7SL1. Two barcoded exogenous 7SL1 sequences were synthesized (Supplementary Table S1).

### RNA interference and plasmid transfection

Two sequential knockdowns were performed using 40 nM of small interfering RNAs (siRNAs) custom ordered from Horizon Discovery (Supplementary Table S1), 72 hours and 24 hours prior to harvest. Knockdowns were performed with siLentFect reagent (Bio-Rad, 703362) according to the manufacturer’s specifications. The control siRNA targeted luciferase mRNA. Transient transfection of plasmid containing exogenous 7SL and U1 sequences was performed 18 hours prior to harvest with Lipofectamine 2000 (Thermo Fisher) according to manufacturer’s recommendations.

### qPCR assays

AR17 (Supplementary Table S1)-primed cDNA was amplified using Fast SYBR Green master mix (Thermo Fisher) containing ROX with 1 μM primers targeting RNAs of interest (Supplementary Table S1) in a 10 μL reaction. Technical duplicates were performed for each sample. Reactions underwent 40 cycles of 95°C denaturation and 60°C annealing with the QuantStudio Real-Time PCR system (Thermo Fisher) using the Fast protocol. Primer products were validated by quantification of primer efficiency, melt curve, and visualization by agarose gel electrophoresis. Cq values were determined with QuantStudio analysis software using default detection. Relative levels were quantified using the ΔΔC_t_ method (48).

### RNA immunoprecipitation

Control or TENT2 KO 293 T-REx cells were transfected with plasmid containing exogenous 7SL and U1 sequences 18 hours prior to harvest. Cells were harvested by scraping into ice cold PBS and flash frozen in liquid nitrogen. Cells were resuspended in ice-cold isotonic lysis buffer (50 mM Tris-HCl pH 7.5, 150 mM NaCl, 5 mM EDTA, 0.5% Triton X-100, 80 U/mL RNaseOut (Thermo Fisher), protease inhibitor tablet/10 mL (Roche)) and incubated with rotation for 20 minutes at 4°C. Cellular debris was pelleted at 13,000 *g* for 15 minutes at 4°C. The supernatant was briefly pre-cleared using Protein G beads (Thermo Fisher, 10004D) for 10 minutes at 4°C with rotation. Beads for immunoprecipitation were washed with bead wash buffer (PBS pH 7.4, 0.1% Tween 20) and subsequently incubated with anti-La antibody (Santa Cruz Biotechnology, sc-80656), anti-SRP54 antibody (Invitrogen, MA5-34835), rabbit IgG (Cell Signaling Technology, 2729), or mouse IgG1 (Cell Signaling Technology, 5415) antibody for 1 hour at 4°C with rotation, then washed with bead wash buffer. Sample protein was quantified with a BCA assay (Thermo Fisher). 1 mg of protein per sample was diluted 5-fold with ice-cold detergent dilution buffer (isotonic lysis buffer without Triton X-100) and incubated with antibody-bead complexes for 1 hour at 4°C with rotation. Samples were washed 5 times with ice-cold immunoprecipitation wash buffer (PBS pH 7.4, 0.1% Tween 20, 150 mM NaCl) at 4°C. RNA from input and immunoprecipitation material was extracted with TRIzol (Thermo Fisher) according to the manufacturer’s recommendation. Gene-specific 3’-end sequencing libraries were prepared as described above. Libraries were sequenced on an Illumina MiSeq platform.

### SLAM-seq

Control or TENT2 KO 293 T-REx cell media was treated with 4-thiouridine [s4U] at a final concentration of 100 µM as previously described and incubated for 3 hours, then refreshed with 100 µM s4U-containing media and incubated for an additional 3 hours (49). Cells were harvested by scraping into ice cold PBS and flash frozen in liquid nitrogen. RNA was extracted with TRIzol (Thermo Fisher) according to the manufacturer’s recommendation.

Following RNA extraction, samples were treated with either iodoacetamide (IAA) or dimethylsulfoxide (DMSO) for 15 minutes at 50°C as previously described (49). Gene-specific 3’-end sequencing libraries were prepared as described above. Libraries were sequenced on an Illumina MiSeq platform. Gene-specific alignments were performed using in-house scripts which permitted T>C conversions. The number of conversions for each read was quantified and reads with two or more conversions were subsequently analyzed as nascently-labeled reads (Supplementary Figure S4G).

### Website Referencing

https://pypi.org/project/jla-demultiplexer - for analysis of Tailer-produced taildata files.

## RESULTS

### Global analysis of sncRNA 3’ end dynamics reveals transient and stable post-transcriptional A- and U-tails

To globally investigate the dynamics of human sncRNA 3’-end processing we performed 3’-end sequencing of steady-state and newly transcribed sncRNAs, approximately 100 to 500 nucleotides in length, from human embryonic kidney (HEK) 293TRex cells (Figure 1A). Newly transcribed sncRNAs (hereafter referred to as nascent sncRNAs) were isolated by click-it chemistry after two hours of metabolic labeling with 5-ethynyl-uridine. A two-hour incubation was necessary for achieving sufficient enrichment over steady state sncRNAs and depth of sequencing (Supplementary Figures S1A and S1B). A wide variety of small RNAs were represented in both steady-state and nascent libraries (Supplementary Figure S1C). Since RNAs that undergo 3’-end processing during biogenesis are expected to show differences in 3’ end nucleotide composition in the nascent versus the steady state populations, we compared 3’ ends of individual RNAs in nascent and steady state. We focused the analyses on full-length sncRNAs (terminating at positions -10 to +50 relative to annotated 3’ ends) to avoid confounding effects of sncRNAs that may have been truncated during RNA isolation. We first examined the dynamics of post-transcriptional A- and U-tailing. We hypothesized that sncRNA post-transcriptional processing may be linked to their mechanism of initial 3’ end formation and therefore grouped sncRNAs according to their transcriptional origin as snoRNAs (including box H/ACA snoRNAs, box C/D snoRNAs and scaRNAs), snRNAs (including spliceosomal snRNAs and U3 and U8 snoRNAs) and Pol-III RNAs.

**Figure 1.**
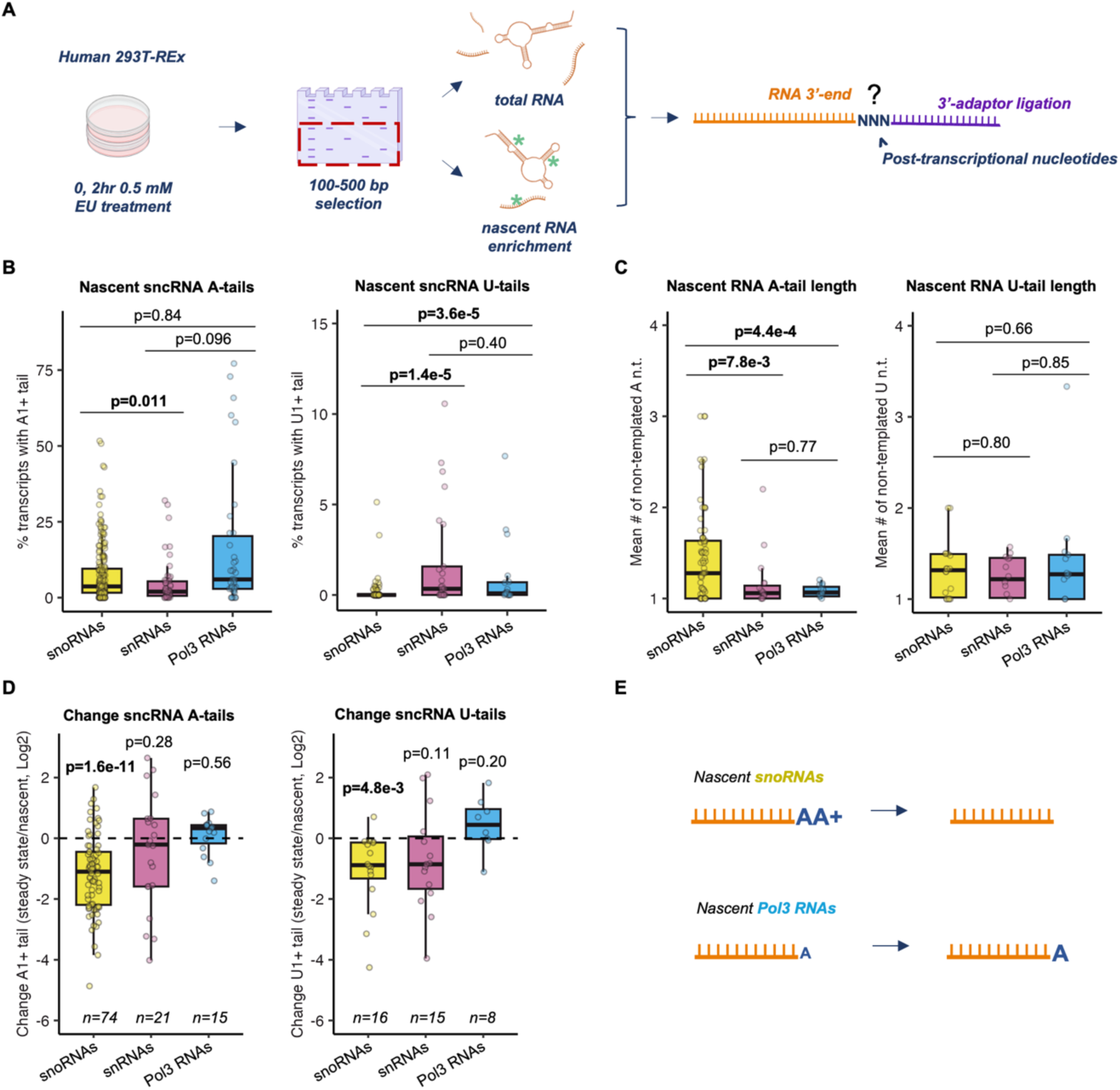
Post-transcriptional A- and U-tails are transient on a subset of sncRNAs but stable on others. **(A)** Schematic of the nascent and steady state sncRNA global 3’ end sequencing workflow. Human embryonic kidney (HEK) 293 T-REx cells were metabolically labeled with 5-ethynyluridine (EU) followed by RNA size-selection and nascent RNA capture. RNA 3’ ends were determined by global RNA 3’ end sequencing of nascent and steady-state sncRNA samples in biological triplicate. Depictions of cell culture, gel extraction, and individual RNAs were created in BioRender (MO27QXAFOF, https://biorender.com/j15w879). **(B)** Percentage of nascent sncRNA species with A- (left) and U-tails (right). RNAs are grouped by their transcriptional origin and dots represent individual sncRNAs. P-values were determined by two-sample Kolmogorov-Smirnov (KS) tests, with p<0.05 shown in bold (snoRNAs n=98, snRNAs n=25, Pol3 RNAs n=19). **(C)** Mean lengths of nascent sncRNA A- (left, snoRNAs n=74, snRNAs n=19, Pol3 RNAs n=15) and U-tails (right, snoRNAs n=18, snRNAs n=14, Pol3 RNAs n=13). P-values were determined by two-sample KS tests, with p<0.05 shown in bold. **(D)** Log2-fold ratios of percentages of A- (left) and U-tailed (right) sncRNA species in steady-state over nascent conditions. Transcripts with 0.1% or greater tailing in both nascent and steady state conditions were plotted. P-values were determined by a one-sample two-tailed t-test against mu=0, with p<0.05 shown in bold. **(E)** Schematic summarizing the 3’-end tail status of nascently-made snoRNAs and Pol-III RNAs. snoRNAs are modified with short oligo(A) tails which are absent in the steady state, while Pol-III RNAs retain their predominant mono(A) tails in the steady state. Depiction of RNA was created in BioRender (MO27QXAFOF, https://biorender.com/j15w879).

Post-transcriptional A-tailing was prevalent among nascent sncRNA species, particularly among Pol-III RNAs, some of which saw A-tailing of over 50% of the nascent population (Figure 1B). SnRNAs and Pol-III RNAs that saw A-tailing were typically mono(A)-tailed, whereas, consistent with previous observations (31, 33), snoRNAs experienced significantly longer oligo(A) tailing (Figure 1C). Post-transcriptional uridylation was also observed for a subset of sncRNAs but at a much lower frequency than A-tailing, and was significantly less frequent among snoRNAs than among snRNAs and Pol-III RNAs (Figures 1B and 1C). Of note, post-transcriptional uridylation of Pol-III RNAs can be difficult to detect given their oligo-uridine termination sequences which cannot be distinguished from post-transcriptional uridines unless U-tails are very long (see further analyses below). Post-transcriptional cytosine and guanosine tails were detectable at some sncRNAs but were very rare and were not further investigated here (Supplementary Figures S1D-F).

To monitor the dynamics of sncRNA A- and U-tailing, we next compared A- and U-tails in steady state versus nascent conditions. This revealed different dynamics for sncRNA species from different transcriptional origins. For snoRNAs, post-transcriptional A- and U-tailing was overwhelmingly transient, as evidenced by a significant shortening of A- and U-tails in steady state compared to nascent conditions (Figure 1D and 1E). By contrast, post-transcriptional A- and U-tails were generally stable for Pol-III RNAs, while snRNAs displayed more of a mixture of transient and stable tails dependent on the RNA species (Supplementary Table S3).

### Transient A-tailing of snoRNAs is associated with instability

Post-transcriptional A- and U-tailing has previously been associated with promotion of RNA 3’ end trimming (1–3, 30, 31, 50) or RNA degradation (3, 50–52) but the principles that dictate one fate over another remain unclear. To first analyze whether A- or U-tailing correlates with 3’ end trimming for each class of sncRNAs, we assessed sncRNA 3’ end trimming by monitoring changes in 3’-ends of RNAs in the steady state population compared to the nascent. As expected, we observed significant 3’ end shortening of a majority of snRNAs and snoRNAs, while a majority of Pol-III sncRNAs saw little 3’-end trimming overall (Figure 2A and Supplementary Figure S2A). This trimming was not restricted to removal of post-transcriptional tails as it generally extended into the genome-encoded sequence (Figure 2B and Supplementary Figure S2B). If 3’ end trimming is promoted by A- or U-tailing it is predicted that RNAs that show 3’ trimming also show A- or U-tailing that is transient. To test this prediction, we classified RNAs with transient A- or U-tailing as those that displayed a significantly (p<0.05 by t-test) higher fraction of A- or U-tailed molecules in the nascent population as compared to steady state. This analysis revealed no significant difference between the 3’ end trimming of transiently A- or U-tailed sncRNAs and the sncRNA population as a whole (Figure 2C and Supplementary Figure S2C).

**Figure 2.**
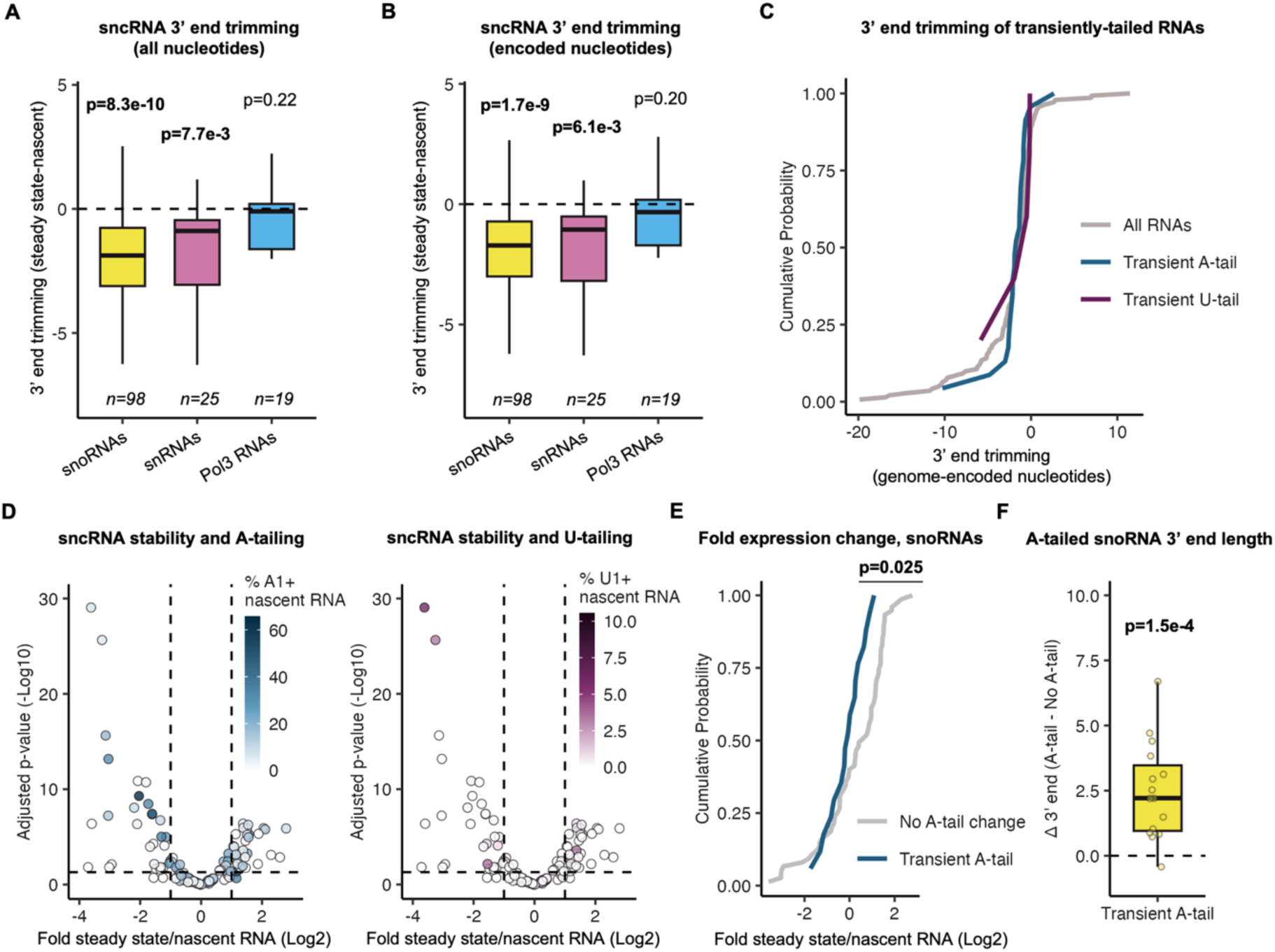
Transient A-tailing is associated with destabilization of snoRNAs. **(A)** and **(B)** Box plots showing ranges of sncRNA 3’ trimming as measured by the difference in the mean 3’-end positions of transcripts in steady-state versus nascent RNA populations. Transcripts were binned by their transcriptional origin. Values in panel *B* excludes nucleotides from post-transcriptional tails. P-values were determined by a one-sample two-tailed t-test against mu=0, with p<0.05 shown in bold. **(C)** Cumulative plot showing sncRNA 3’ trimming as measured by the difference in the mean 3’-end positions of transcripts in steady-state versus nascent RNA populations. SncRNAs that saw transient 3’ A-or U-tailing, as measured by a significantly (p<0.05; two-sample two-tailed t-test) higher fraction of A- or U-tailed molecules in nascent over steady state populations, are compared to all RNAs (All RNAs n=142, transient A-tails n=23, transient U-tails n=5). **(D)** Volcano plot comparing sncRNA stability with the percentages of A- (left) or U-tailed (right) molecules. The log2 fold ratio of sncRNA levels in steady-state over nascent conditions was quantified with DESeq2 and plotted against -log10-converted adjusted p-values. Individual sncRNAs are colored according to their percentage of modification with A- or U-tails of any length. The horizontal dashed line represents p=0.05. **(E)** Cumulative plot showing snoRNA stability as measured by the log2 ratio of steady-state over nascent levels. snoRNAs with significantly higher A-tailing in the nascent population relative to steady state are compared to all other snoRNAs. P-value was determined by a two-sample KS test (no A-tail change n=75, transient A-tail n=17). **(F)** Box plot showing the mean position of snoRNA A-tails relative to the 3’ end position of their unadenylated counterparts for transiently A-tailed snoRNAs. P-value was determined by a one-sample two-tailed t-test against mu=0 (n=15).

We next assessed whether a correlation exists between 3’-end tailing and sncRNA stability. RNAs that are unstable are expected to be enriched in the nascent population over steady state. We therefore used the ratio of RNA abundance in the steady state over nascent population as a proxy for RNA stability. Multiple sncRNAs that are unstable by this measure showed high levels of A- or U-tailing in the nascent population (Figure 2D). Evaluating whether a correlation exists between transient tailing and stability for individual groups of sncRNAs, in the case of snoRNAs, those that underwent transient A-tailing showed significantly lower steady state to nascent RNA ratios than the remainder of the snoRNA population suggesting that the transiently A-tailed population is unstable (Figure 2E). Consistent with previous observations by others (31, 53, 54), the transient snoRNA A-tails were found on snoRNAs not fully processed at their 3’-ends (Figure 2F), suggesting that oligo(A)-tailing and degradation targets snoRNAs that experience stalling in 3’ end processing. In contrast to a previous study suggesting that oligoadenylation is specific to H/ACA box snoRNAs (31), we observed transient oligoadenylation of snoRNAs of all types (Supplementary Table S4). Other classes of RNAs showed very few transiently A-tailed species and no evidence of associated instability (Supplementary Figure S2D). While there were too few transiently U-tailed sncRNAs to be analyzed in a similar manner, the few sncRNAs that were uridylated at 2.5% or greater ratios in the nascent population were, as a group, significantly less stable than other sncRNAs (Supplementary Figure S2E).

### A majority of Pol-III RNA species see subpopulations with stable mono(A)-tailing

We next turned our attention to the short post-transcriptional tails observed on Pol-III RNAs and snRNAs. In contrast to snoRNAs, which typically see transient A-tails, over half of observed Pol-III RNA species, as well as a smaller subset of snRNA species, accumulate post-transcriptional A-tails at significantly higher levels at steady-state than in the nascent population suggesting that these tails are stable (Figure 1D and 3A). Pol-III RNAs as a group accumulated with A-tails at steady state at a higher level than other sncRNAs (Figure 3B) and these A-tails were generally mono(A)-tails (Figure 3C). While a majority of Pol-III RNA species can be observed with post-transcriptional A-tails at steady state, only three (7SL1, 7SL2 and 7SL3 RNAs) accumulate with A-tails on more than half of their population (Figure 3D and Supplementary Table S5). Performing the same analysis for U-tailing revealed a subset of snRNAs that accumulate U tails in the steady state, as well as a select few snRNAs that see U-tails that are transient (Supplementary Figure S3).

**Figure 3.**
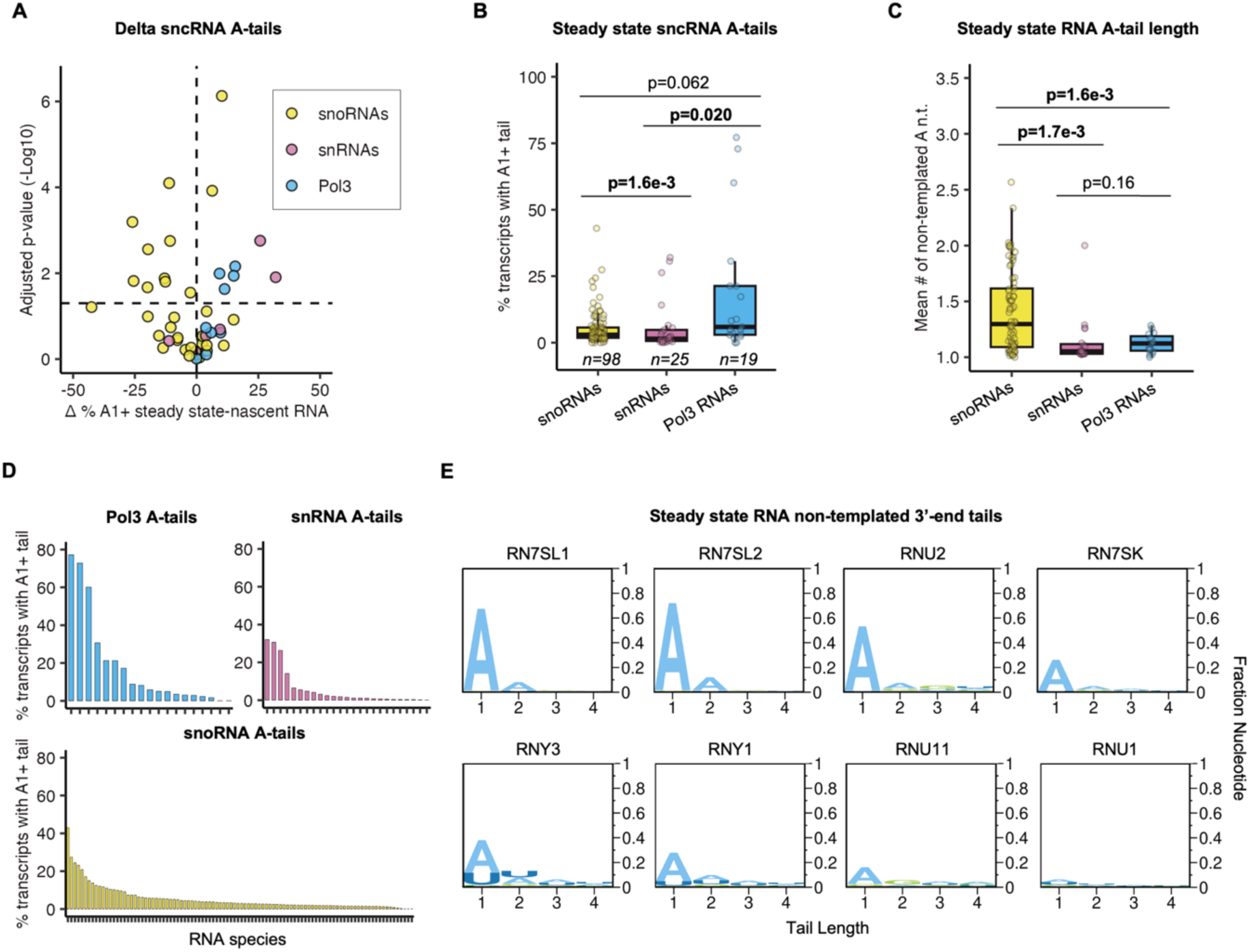
A subset of Pol-III RNAs and snRNAs stably accumulate with mono-A tails. **(A)** Volcano plot showing differences in percentages of A-tailed sncRNAs in the steady-state versus nascent conditions versus the -Log10 p-value of the difference. Only sncRNAs with ≥5% of the population with A tails in the steady state are plotted. Horizontal line indicates p=0.05 by a two-sample two-tailed t-test. Transcripts are colored by their transcriptional origin. **(B)** Box plots showing ranges in percentages of sncRNAs with A-tails at steady state. Transcripts were binned by their transcriptional origin. P-values determined by two-sample KS tests, with p<0.05 shown in bold. **(C)** The mean length of steady state A-tails of transcripts. Transcripts are binned by their transcriptional origin. P-values were determined by two-sample KS test, with p<0.05 shown in bold (snoRNAs n=74, snRNAs n=19, Pol3 RNAs n=15). **(D)** Rank order plot of sncRNAs by the percentage of the population with A-tails at steady state. Transcripts were plotted separately by their transcriptional origin. **(E)** Logo plots showing post-transcriptional tails at steady state for directly sequenced Pol-III RNAs and snRNAs. Post-transcriptional nucleotides are plotted as fractions of the total population.

To confirm the mono(A)-tailing observed in our global 3’-end sequencing data, we monitored 3’ ends of a subset of Pol-III RNAs and snRNAs by gene-specific 3’ end sequencing, with U1 snRNA serving as a mostly unadenylated control. Plotting the composition of post-transcriptional tails for these RNAs support the conclusion that these RNAs see 3’ monoadenylation at the steady state, ranging from ≈15% of the population for U11 snRNA to ≈70% for 7SL1 and 7SL2 RNAs (Figure 3E). These levels of monoadenylation are consistent with previous observations for individually tested Pol-III RNAs and snRNAs (33). As expected, the U1 snRNA negative control showed little post-transcriptional 3’-end tailing at steady state. These observations, taken together, demonstrate that mono(A) tails stably accumulate to the steady state on a subset of the population of a majority of Pol-III RNA species as well as a subset of snRNAs.

### The non-canonical polymerase Terminal Nucleotidyltransferase 2 (TENT2) promotes Pol-III and snRNA 3’ end mono(A)-tailing

To better understand how 3’-end monoadenylation may impact sncRNA processing or function, we sought to identify the enzymatic activity responsible for this modification. The non-canonical polymerases TENT4A and TENT4B have been previously observed to adenylate a large number of sncRNAs (2, 31, 50, 51, 55). To assess whether TENT4A/4B are responsible for the observed oligo(A)- and/or mono(A)-tailing of sncRNAs, we analyzed a published TENT4A/4B knockdown 3’-end RNA-seq dataset (56). Plotting the changes in sncRNA 3’-adenylation in control versus TENT4A/4B co-depletion conditions revealed that snoRNA adenylation was significantly decreased during TENT4A/4B knockdown (Figure 4A and Supplementary Figure S4A). This is consistent with previous reports of snoRNA adenylation by TENT4B (2, 31). By contrast, snRNA and Pol-III RNA A-tailing was not significantly impacted. Taken together, these observations implicate TENT4A/4B in the transient oligo(A)-tailing of snoRNAs, while the short A-tails observed on Pol-III RNAs and a subset of snRNAs appear to be added by different polymerase(s).

**Figure 4.**
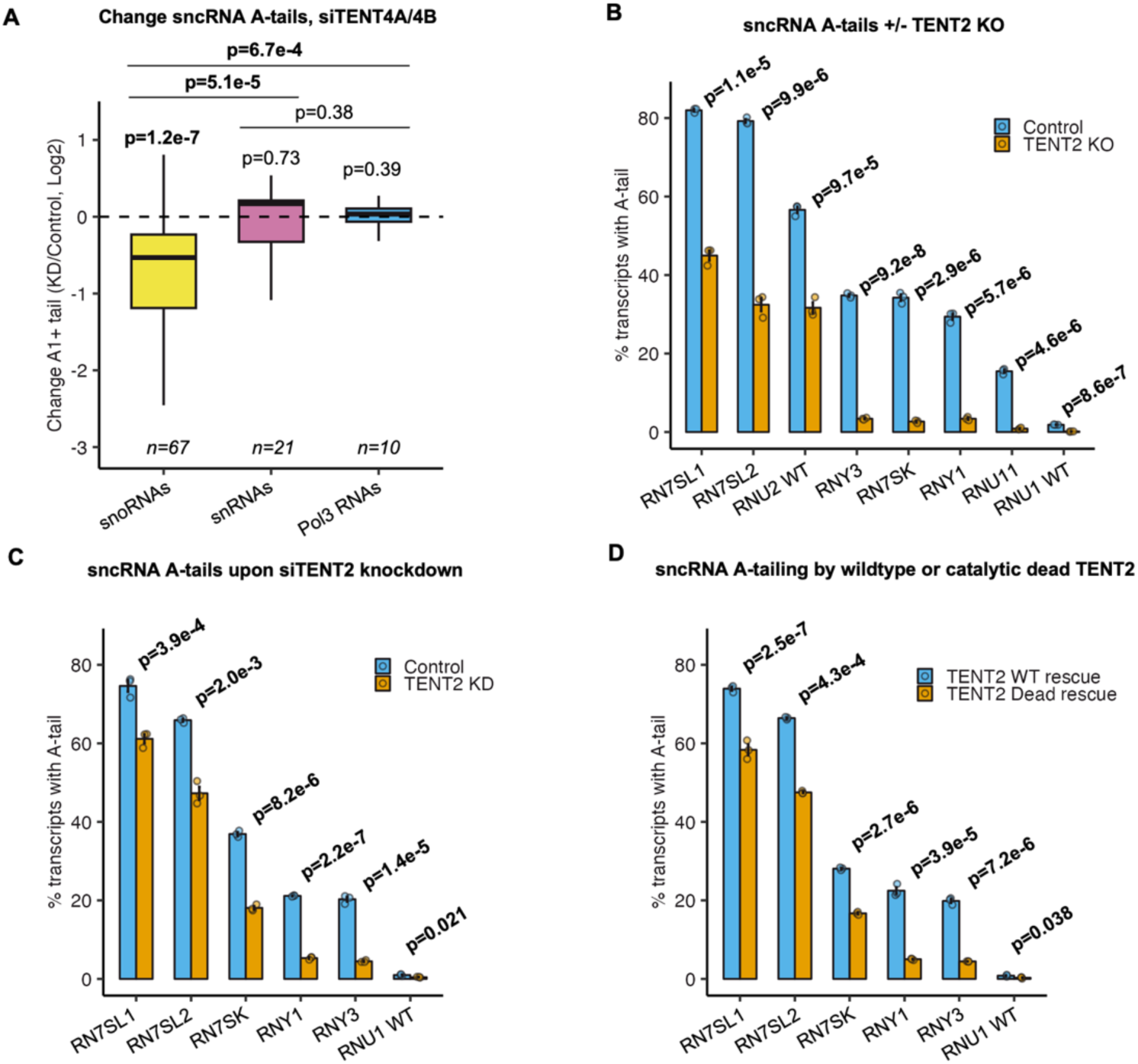
TENT2 contributes to Pol3 and snRNA 3’ end A-tailing. **(A)** Box plots showing Log2 ratios in percentages of A-tailed sncRNAs in TENT4A/4B knockdown conditions over to control conditions. The dataset is from (Lim 2018). Transcripts are grouped by their transcriptional origin. P-values for individual groups were determined by one-sample two-tailed t-tests against mu=0. P-values between groups were determined by two-sample KS tests. p<0.05 is shown in bold. **(B)** Percentage of adenylated sncRNAs in TENT2 KO or control conditions in HEK 293T-Rex cells as measured for select Pol-III RNAs and snRNAs by gene-specific 3’ end sequencing. Data is represented as mean +/- standard error of the mean (SEM) and p-values were determined by two-sample two-tailed t-tests, with p<0.05 shown in bold (n=3 for each transcript). **(C)** and **(D)** Percentage of adenylated sncRNAs in TENT2 siRNA depletion or control conditions in HEK 293T-Rex cells (panel *C*) or in TENT2-depleted HEK 293T-Rex cells complemented with TENT2 wild-type or TENT2 catalytic dead proteins (panel *D*). Data is represented as mean +/- SEM and p-values were determined by two-sample two-tailed t-tests, with p<0.05 shown in bold (n=3 for each transcript).

Previous studies identified the mouse protein Germ Line Development 2 (mGLD2) as an enzyme that participates in monoadenylation of 7SL RNA (29). To test whether the human homolog of mGLD2, TENT2, promotes sncRNA 3’-monoadenylation, we knocked out the *TENT2* gene in HEK 293T-REx cells. Following confirmation of *TENT2* knockout (Supplementary Figures S4B and S4C), we sequenced the 3’ end of the subset of Pol-III RNAs and snRNAs that presented with at least 20% of the population with post-transcriptional adenylation at steady state. This revealed a significant reduction in 3’-adenylation for all RNAs tested (Figure 4B and Supplementary Figure S4D). Adenylation was almost completely abolished for 7SK, Y1, Y3 and U11 RNAs, whereas for 7SL1, 7SL2 and U2 RNAs approximately half of the original 3’-adenylated population remained after TENT2 knockout. The impact of TENT2 on Pol-III RNA and snRNA adenylation was confirmed by siRNA-mediated depletion of TENT2 (Figure 4C and Supplementary Figure S4E), and by add-back of exogenous wild-type TENT2 versus catalytically inactive TENT2 (Figure 4D). 3’ end sequencing of nascent 7SL and U2 RNAs isolated from TENT2 knockout and control cells revealed a similar impact on 3’-adenylation as observed at steady state, showing that the adenylated population resistant to TENT2 KO does not reflect a highly stable population generated prior to TENT2 KO (Supplementary Figures S4F and S4G). Thus, snRNAs and Pol-III RNAs are mono-adenylated by TENT2, with a subset of the 7SL and U2 RNA population adenylated by an additional unknown enzyme, which is not TENT4A/4B (Supplementary Figure S4H).

### Monoadenylation by TENT2 inhibits trimming and extension of Pol-III RNA 3’ uridine tails

Adenylation has previously been observed to inhibit uridylation of tested Pol-III RNAs *in vitro* and when injected into nuclei of *Xenopus* oocytes (34). We therefore examined the impact of TENT2 on cellular Pol-III RNA uridylation and deuridylation dynamics. While U6 snRNA is known to undergo uridylation during biogenesis, many other Pol-III RNAs have instead been observed to undergo various levels of trimming of their genome-encoded uridine tails (15–18, 57). Indeed, comparing Pol-III RNA 3’ end uridine termini in our 3’ end sequencing data for nascent and steady-state populations revealed that while U6 snRNA undergoes 3’ uridine extension during biogenesis, a majority of Pol-III RNAs experience significant trimming of 3’ uridines (Figure 5A and Supplementary Table S6). To test whether mono(A)-tailing by TENT2 impacts Pol-III RNA 3’ end trimming, we used direct sequencing to monitor the effect of TENT2 knockout on the mean number of 3’-uridines of select Pol-III RNAs. This revealed a general reduction in the number of 3’ end uridines of the tested Pol-III RNAs upon TENT2 KO, with the most highly adenylated Pol-III RNAs, 7SL1 and 7SL2 RNAs, showing significant shortening (Figure 5B). Thus, monoadenylation by TENT2 stalls 3’ uridine trimming of 7SL RNAs and possibly other Pol-III RNAs.

**Figure 5.**
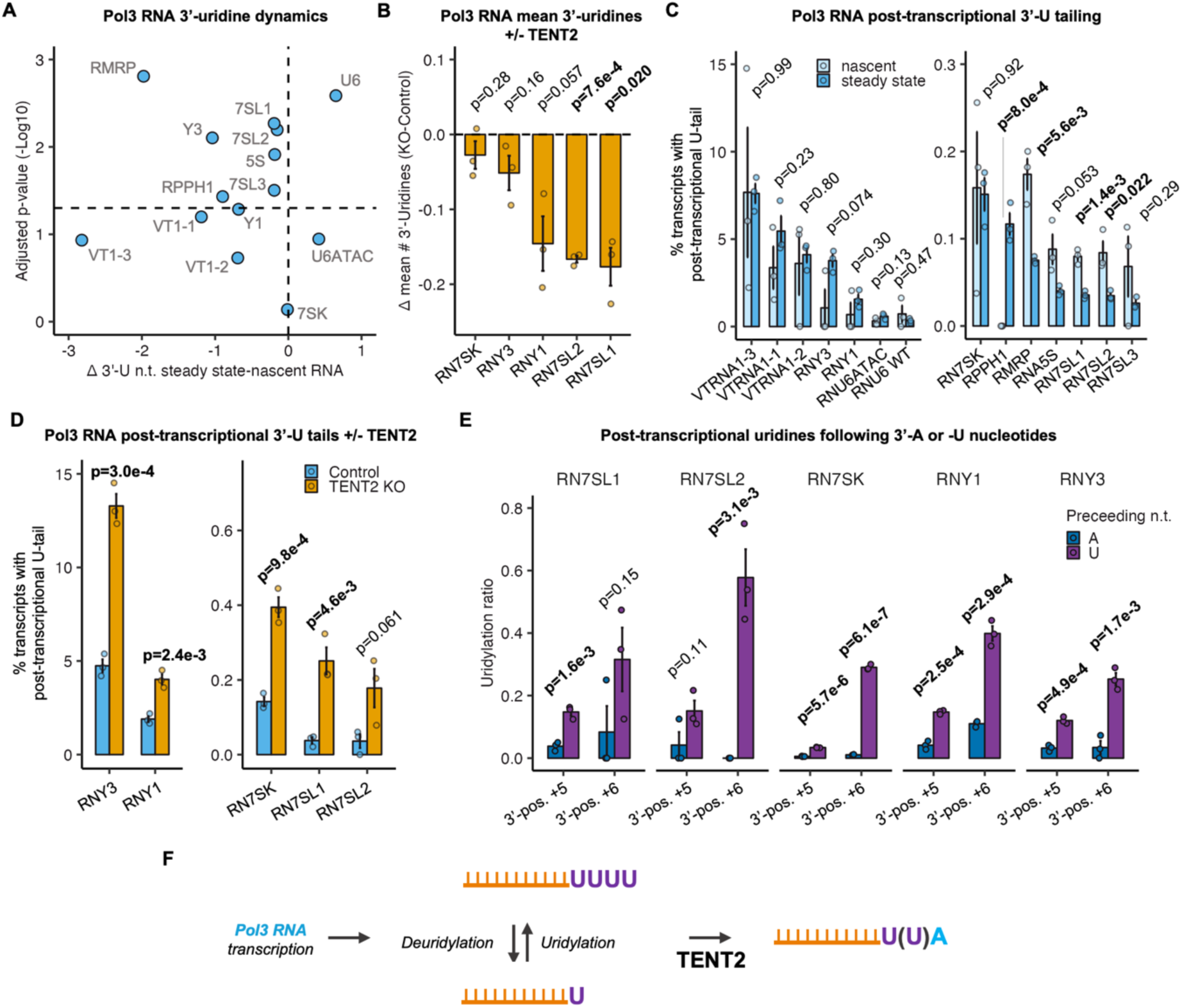
Mono(A) tails inhibit trimming and tailing of Pol3 RNA oligo(U)-termini. **(A)** Differences in lengths of 3’ uridine termini of steady state versus nascent Pol-III RNAs plotted against the -Log10 p-value for three independent experiments. P-values were determined by two-sample two-tailed t-tests. The horizontal line represents p=0.05 (n=3 for each transcript). **(B)** Difference in mean number of 3’-uridines of Pol-III RNAs between TENT2 KO and control cells. Data is represented as mean +/- SEM and p-values were determined by one-sample two-tailed t-tests against mu=0 with p<0.05 shown in bold (n=3 for each transcript). **(C)** Percentage of Pol-III transcripts with post-transcriptional 3’-U tails in nascent or steady state conditions. Only uridine(s) following the final uridine of their termination sequence are plotted. A termination sequence length of 4 was used for 7SL1-3, 7SK, Y1, Y3, VTRNA1-1, VTRNA1-2, and VTRNA1-3. A termination length of 5 was used for U6, 5S, and RPPH1. A termination length of 6 was used for MRP. A termination length of 13 was used for U6ATAC. Data is represented as mean +/- SEM and p-values were determined by two-sample two-tailed t-test, with p<0.05 shown in bold (n=3 for each transcript). **(D)** Percentage of Pol-III transcripts with post-transcriptional 3’-U tails in control or TENT2 KO conditions. Data is represented as mean +/- SEM and p-values were determined by two-sample two-tailed t-test, with p<0.05 shown in bold (n=3 for each transcript). **(E)** Ratio of post-transcriptional U nucleotides preceded by either A or U nucleotides plotted for 3’-end positions +5 and +6 relative to the first nucleotide of the termination sequence (defined as +1) so as to ensure a post-transcriptional origin of the uridine. Data is represented as mean +/- SEM and p-values were determined by two-sample two-tailed t-tests, with p<0.05 shown in bold (n=3 for each transcript). **(F)** Schematic representing uridine extension and trimming of nascent Pol-III RNAs which is terminated by monoadenylation by TENT2. Depiction of RNA was created in BioRender (MO27QXAFOF, https://biorender.com/j15w879).

Further analyses of our genome-wide sequencing data also revealed that all Pol-III RNA species see a fraction of their populations with post-transcriptional uridylation beyond their genome-encoded U-tails (Figure 5C). For a subset of Pol-III RNAs, including 7SL1 and 7SL2 RNAs, this post-transcriptional uridylation was significantly more prevalent in the nascent than the steady-state population, suggesting that it is associated with biogenesis. The post-transcriptionally uridylated populations increased upon TENT2 knockout for each of the Pol-III RNA species monitored by direct sequencing (Figure 5D), suggesting that mono(A)-tailing inhibits post-transcriptional uridylation. Consistent with this idea, post-transcriptional uridines were preceded by uridines significantly more often than by adenosines for each of these RNAs (Figure 5E). Taken together, these observations show that mono(A)-tailing by TENT2 inhibits both 3’-uridine trimming and extension of Pol-III RNAs (Figure 5F).

### TENT2 inhibits 7SL RNA association with La protein

Given that 7SL1 and 7SL2 RNAs are the most highly adenylated sncRNAs with over 70% of the population accumulating with mono-A-tails, we next asked whether mono(A)-tailing by TENT2 impacts 7SL RNA biogenesis. We first asked if mono(A)-tailing by TENT2 affects 7SL RNA accumulation. Examining 7SL RNA levels in the presence or absence of TENT2 showed no effect of TENT2 depletion on 7SL RNA steady state accumulation (Supplementary Figure S5A). Given that 7SL RNAs are highly stable molecules we also monitored the impact of TENT2 on the accumulation of transiently expressed exogenous 7SL1 RNAs. The exogenous 7SL1 RNAs, for reasons that are unclear, are less adenylated and more uridylated at the 3’ end than endogenous 7SL RNAs and more dependent on TENT2 for their 3’ adenylation (Supplementary Figures S5B-E). We therefore wondered whether the exogenous 7SL1 RNAs would be more dependent on TENT2 for their accumulation. However, TENT2 depletion did not impact the accumulation of exogenous 7SL1 RNAs as monitored by RNA sequencing (Supplementary Figure S5F). These observations suggest that TENT2 does not significantly impact 7SL RNA stability.

Another possible impact of 7SL RNA monoadenylation could be on its association with RNA-binding proteins. The abundant nuclear RNA binding protein La is known to have high affinity for RNAs with 3’-oligouridines (58–60) and has been previously observed to associate with 7SL RNA (61, 62). Performing immunoprecipitation against La and assessing associated 7SL RNA levels in the presence or absence of TENT2 revealed significantly increased 7SL RNA association with La in the absence of TENT2 (Figure 6A). Sequencing the 3’ ends of the La-associated 7SL RNAs revealed a significant enrichment in 3’ uridylated and de-enrichment in adenylated species as compared with the overall 7SL RNA population (Figure 6B), consistent with a preference of La for unadenylated 7SL RNAs. Moreover, exogenous 7SL1 RNAs were enriched in association with La over the endogenous 7SL RNAs (Figure 6C), consistent with their significantly lower levels of adenylation and higher levels of uridylation (Figure 6D and Supplementary Figure S5E). These observations taken together demonstrate that mono(A)-tailing by TENT2 terminates 3’ uridine trimming at 7SL RNA 3’ ends and inhibits the association of 7SL RNAs with La protein, consistent with the known preference of La for RNAs with oligo-uridine 3’-termini. Other tested sncRNAs were not observed to be significantly enriched with La upon TENT2 depletion (Supplementary Figure S5G), perhaps reflecting the smaller fractions of these RNA populations that receive mono(A)-tails by TENT2.

**Figure 6.**
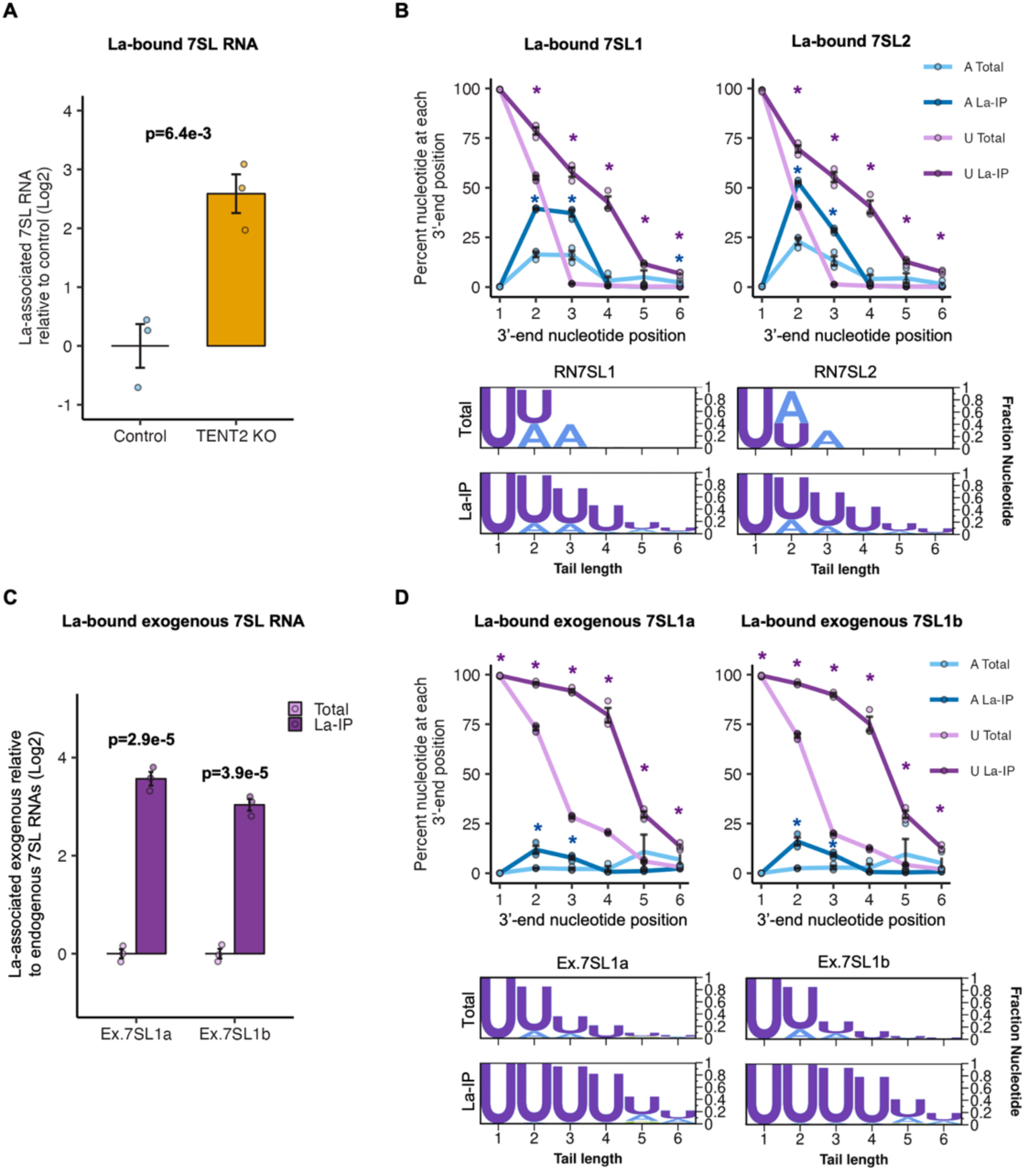
7SL RNA association with La protein is inhibited by TENT2. **(A)** Levels of 7SL RNAs associated with La in control versus TENT2 KO conditions monitored by IP followed by RT-qPCR for 7SL RNAs relative to U1 snRNAs and normalized against the IgG IP controls. Data is represented as mean +/- SEM and p-value was determined by a two-sample two-tailed t-test, with p<0.05 indicated in bold (n=3 for each condition). **(B)** 3’ end nucleotide compositions of steady-state versus La-associated endogenous 7SL RNAs shown as line plots (top) and logo plots (bottom). Nucleotide positions are shown relative to the first nucleotide of the 7SL termination sequence, defined as position +1. Data is represented as mean +/- SEM and p-values between IP and input groups were determined by two-sample two-tailed t-tests, with p<0.05 indicated by asterisks (n=3 for each condition). Purple lines and asterisks represent U-tails, and blue lines/asterisks represent A-tails. **(C)** Levels of exogenous 7SL1a and 7SL1b RNAs associated with La relative to endogenous 7SL RNAs monitored by IP followed by sequencing. The relative abundance of exogenous 7SL1 RNAs bound to La was compared to the mean of endogenous 7SL1 and 7SL2 RNAs, then normalized to the relative abundance of exogenous U1 to endogenous U1. Data is represented as mean +/- SEM and p-values were determined by one-sample two-tailed t-tests against mu=0 with p<0.05 indicated in bold (n=3 for each condition). **(D)** Same analysis as panel *B*, but monitoring exogenous 7SL1 RNAs.

### Excessively uridylated 7SL RNAs are impaired in the final step of SRP assembly

La-associated RNAs have been reported to be retained in the nucleus (15, 58, 63–65). Given that the last step of SRP assembly, the association of 7SL RNA with SRP54, occurs in the cytoplasm we considered the possibility that association of 7SL RNA with La negatively impacts its subsequent nuclear export and assembly with SRP54 (37–39). Given the preference of La for RNAs with three or more 3’ terminal uridines (58–60), we tested the prediction that 7SL RNAs with three or more 3’ end uridines should be de-enriched in their association with SRP54. Indeed, immunoprecipitation against SRP54 followed by 7SL RNA 3’ end sequencing revealed a significant de-enrichment of this population of molecules in association with SRP54 for both 7SL1 and 7SL2 RNAs (Figure 7A). This could be observed both in the presence or absence of TENT2 KO. A similar de-enrichment in association with SRP54 was observed for exogenous 7SL1 RNAs with three or more 3’ uridines (Figure 7B), and, consistent with their significantly higher levels of uridylation, exogenous 7SL1 RNAs showed significantly lower levels of association with SRP54 than endogenous 7SL RNAs (Figure 7C). The overall level of endogenous 7SL RNA associated with SRP54 was not significantly impacted upon TENT2 KO (Figures 7D) as may be expected given the low fraction (<2%) of endogenous 7SL RNAs that contain three or more 3’ uridines at steady-state (Figure 7A and Supplementary Figure S6A). However, the more extensively uridylated exogenous 7SL1 RNAs were de-enriched in association with SRP54 upon TENT2 KO (Figure 7D) and SRP54-associated exogenous 7SL RNAs could be observed to be significantly de-enriched for uridylated species (Supplementary Figure S6B). Taken together, these observations suggest that monoadenylation by TENT2 promotes 7SL RNA biogenesis by preventing the association of 7SL RNAs with La, which in turn allows 7SL RNAs to assemble with SRP54 in the cytoplasm to complete SRP biogenesis (Figure 7E).

**Figure 7.**
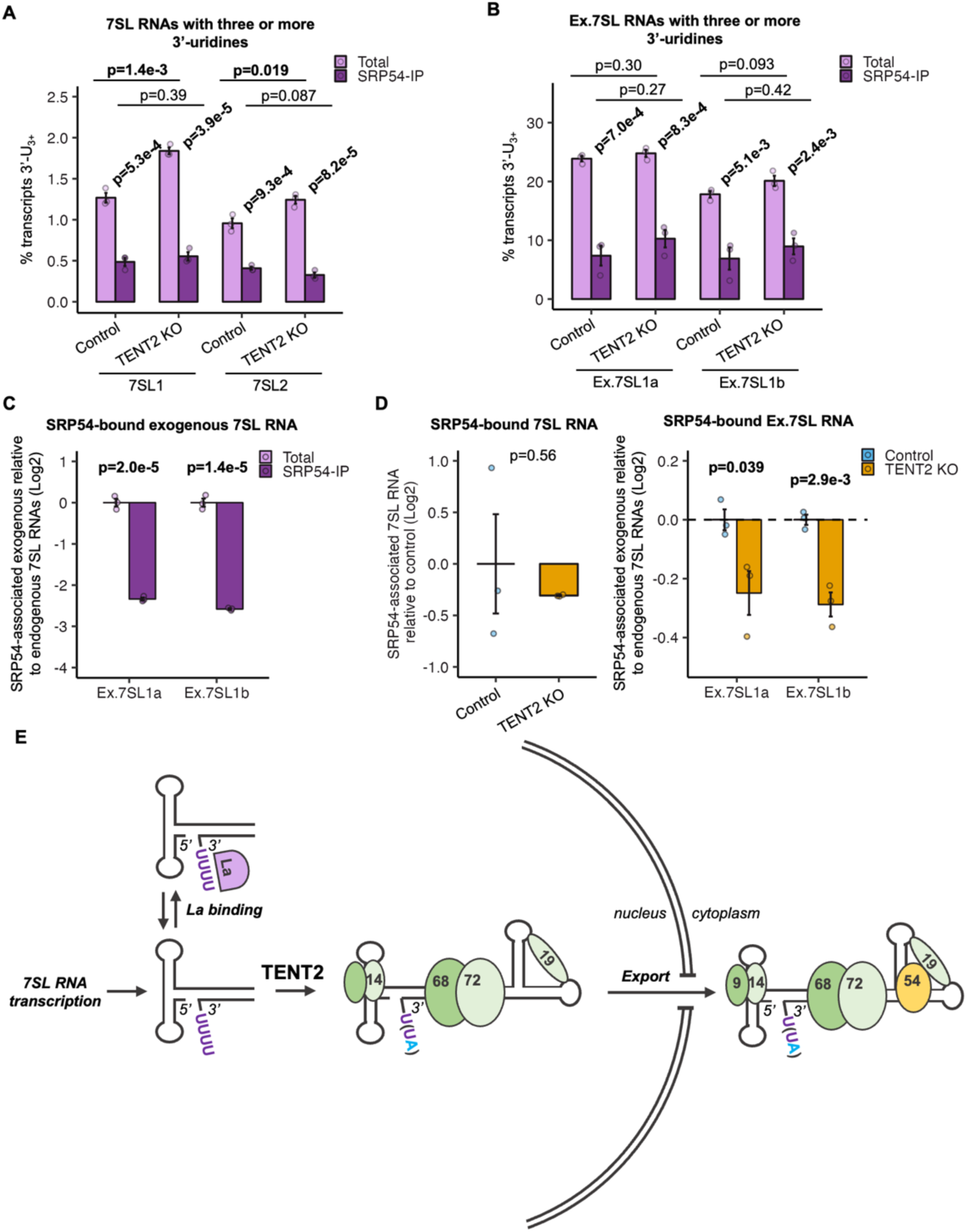
Excessively uridylated 7SL RNAs are impaired in the final step of SRP assembly. **(A)** Percentage of endogenous 7SL RNAs with 3 or more 3’-uridines in input versus SRP54-IP conditions and control versus TENT2 KO conditions. Data is represented as mean +/- SEM and p-values were determined by two-sample two-tailed t-tests, with p<0.05 indicated in bold (n=3 per condition). **(B)** Same as panel *A* but monitoring exogenous 7SL RNAs. **(C)** Log2 ratio of SRP54-associated exogenous 7SL1 RNAs over endogenous 7SL RNAs, monitored by sequencing. Data is represented as mean +/- SEM and p-values were determined by one-sample two-tailed t-tests against mu=0 with p<0.05 indicated in bold (n=3 for each condition). **(D)** Impact of TENT2 KO on 7SL RNA association with SRP54. Left: levels of SRP54-associated 7SL RNAs monitored by IP followed by RT-qPCR for 7SL RNAs relative to U1 snRNAs and normalized against the average level of association in the absence of TENT2 KO (control). Data is represented as the log2 mean +/- SEM and p-value was determined by a two-sample two-tailed t-test, with p<0.05 indicated in bold (n=3 per condition). Right: log2 ratio of exogenous 7SL1 RNAs over endogenous 7SL RNAs in SRP54 IP samples in control and TENT2 KO conditions monitored by sequencing and normalized against the average association ratio in the absence of TENT2 KO (control). Data is represented as the log2 mean +/- SEM and p-values were determined by two-sample two-tailed t-tests, with p<0.05 indicated in bold (n=3 for each condition). **(E)** Schematic representing stages of 7SL biogenesis. 7SL RNAs with 3 or more uridines may be bound by La. TENT2 adenylates 7SL 3’ ends which promotes release from La, allowing nuclear export and assembly of SRP in the cytoplasm with SRP54.

## DISCUSSION

In this study we globally characterized the dynamics of human sncRNA 3’-end processing and the relation between post-transcriptional tailing and sncRNA maturation and degradation. We observed post-transcriptional A-tailing of nascent sncRNAs to be wide-spread and consist of two major types. One is characterized by transient oligo(A)-tailing, which was observed predominantly on nascent snoRNAs and scaRNAs that were not fully processed at their 3’ ends (Figure 1 and Supplementary Figure S2D). This oligo(A)-tailing is carried out by non-canonical polymerases TENT4A/4B and associated with instability rather than maturation (Figure 2), consistent with the known link between TENT4A/4B and the nuclear exosome (50, 51, 55, 66, 67). The second type of sncRNA A-tailing is characterized by mono(A)-tailing, which occurs on a majority of Pol-III RNAs and a smaller subset of snRNAs, and, in contrast to oligo(A)-tailing, stably accumulates to the steady state (Figure 3).

Mono(A)-tailing is broadly carried out by TENT2, although other polymerases appear to contribute as well (Figure 4), and it inhibits Pol-III RNA 3’ uridine trimming and extension (Figure 5). Mono(A)-tailing of 7SL RNAs inhibits their interaction with La protein (Figure 6) and extensively uridylated 7SL RNAs, which are favored for La binding, are inhibited from assembling with the cytoplasmic SRP component SRP54 (Figure 7), suggesting that mono(A)-tailing of 7SL RNAs promotes proper SRP biogenesis.

Our observation of transient oligo(A)-tailing of snoRNAs that are not fully 3’ end processed is consistent with previous evidence for snoRNA adenylation by TENT4B occurring in competition with deadenylation by the deadenylase PARN during snoRNA maturation (2, 3, 27). We observed that the transient oligo(A)-tailing of snoRNAs correlates with instability rather than maturation (Figure 2E), which, together with the observation that tailing occurs primarily on snoRNAs that are not fully 3’ end processed (Figure 2F), suggests that adenylation by TENT4A/4B promotes degradation of snoRNAs whose maturation have stalled during 3’ end processing. We observed transient A-tailing and associated instability for H/ACA and C/D box snoRNAs and scaRNAs alike, suggesting that this mechanism of regulation is broadly applicable to these RNAs (Supplementary Table S4). These observations suggest that the competition between adenylation by TENT4A/4B and deadenylation by PARN (2, 31) at snoRNA 3’-ends dictates whether these RNAs are processed to maturation or subjected to degradation. A similar mechanism has been observed to regulate telomerase RNA levels, whereby deadenylation by PARN protects TERT from degradation initiated by TENT4B-mediated oligoadenylation (51). Similar oligoadenylation/deadenylation dynamics have additionally been observed for miRNAs (68, 69) and snRNAs (1, 3, 4), suggesting that this is a widespread mechanism to control sncRNA expression.

The second type of post-transcriptional A-tailing that we observed consists of mono(A)-tailing, which we find is widespread among Pol-III RNAs and accumulates into the steady state. We identify TENT2 as broadly responsible for this monoadenylation event. Pol-III RNAs naturally terminate with 3’-uridines, and we observed that TENT2 inhibits the trimming of these U-tails that can be observed for a majority of Pol-III RNAs. Thus, mono(A)-tailing by TENT2 may serve to terminate 3’ end processing of Pol-III RNAs, reminiscent of how a 2’,3’ cyclic phosphate modification terminates processing of U6 and U6atac RNAs (32, 70).

Another consequence that we observed of mono(A)-tailing of Pol-III RNAs is inhibition of post-transcriptional uridylation. This is consistent with previous observations of adenylation of Pol-III RNAs inhibiting uridylation *in vitro* and upon injection into *Xenopus* oocyte nuclei (34). TENT2 has previously been observed to stabilize a variety of miRNAs in mice (28, 29). Thus, it is possible that mono(A)-tailing likewise stabilizes Pol-III RNAs via inhibition of uridylation, a known trigger of RNA degradation in the cytoplasm (22, 71). We did not observe evidence for decreased levels of Pol-III RNAs in the absence of TENT2, however, for the majority of Pol-III RNAs, mono(A)-tailing occurs on less than half of the population and it is therefore possible that the mono(A)-tailed population is insufficiently large to observe stabilization of the overall RNA species.

We observed 7SL RNAs as the most highly mono(A)-tailed RNAs with over 70% of the population accumulating with mono(A)-tails at steady state. TENT2 is partly responsible for this adenylation, but our findings suggest that additional polymerase(s) are involved as well. This second polymerase could be PAP-γ which has been previously shown capable of adenylating 7SL RNAs *in vitro*, though RNA specificity for the modification was not observed (72). Our observations that TENT2 depletion leads to increased association of 7SL RNAs with La and that extensively uridylated 7SL RNAs are impaired in assembly with SRP54 suggest that mono(A)-tailing by TENT2 promotes 7SL RNA biogenesis. TENT2 has been reported to localize in the cytoplasm of vertebrates (73) although there has been reports of additional localization in the nucleus in mouse and *Xenopus* (74, 75). Our observations on the impact of TENT2 on 7SL RNA La binding suggests that TENT2 is active in the nucleus of human cells. Given evidence from others that La protein association results in nuclear retention (15, 58, 63–65), the simplest interpretation of our observations is that nascent 7SL RNAs with unprocessed U-tails are retained in the nucleus by La, which prevents them from assembling into mature SRP particles in the cytoplasm. In this scenario, mono(A)-tailing by TENT2 would promote SRP assembly by inhibiting La association and allowing for nuclear export (Figure 7E). An important question for future studies is what drives these 3’ uridine trimming and monoadenylation decisions, and whether these dynamics control the decision of whether Pol-III transcripts are destined for maturation or degradation.

## DATA AVAILABILITY

Global and gene-specific 3’ end RNA sequencing data have been deposited to the Gene Expression Omnibus (GEO) (76) under accession numbers GSE287260 and GSE287259, respectively.

## AUTHOR CONTRIBUTIONS

C.O. and J.L-A. conceptualized the project. C.O. performed all experiments. C.O., B.S., T.N-S. and S.R.L. produced key reagents. C.O., B.S., A.P., T.N-S. and J.L-A. performed data analyses. C.O. and J.L-A. wrote the manuscript.

## ACKNOWLEDGEMENTS

SncRNA global 3’-end sequencing was conducted at the UC San Diego IGM Genomics Center utilizing an Illumina NovaSeq 6000 that was purchased with funding from a National Institutes of Health SIG grant (#S10 OD026929). We thank Kristen Jepsen and the Sanford Human Embryonic Stem Cell Core for assistance with Illumina platform sequencing. We thank Triton Shared Computer Cluster (TSCC) for facilitating computationally-intensive analyses and staff for guidance. We thank Tiantai Ma for his synthesis of exogenous U1 plasmids.

## FUNDING

This work was supported by National Institutes of Health (NIH) grant R35 [GM118069] awarded to J. L.-A.

## CONFLICT OF INTEREST

The authors declare no competing financial interests.

## Notes

https://www.ncbi.nlm.nih.gov/geo/query/acc.cgi?acc=GSE287259

https://www.ncbi.nlm.nih.gov/geo/query/acc.cgi?acc=GSE287260

### Competing Interest Statement

The authors have declared no competing interest.

### Summary of Updates

The title has changed to highlight the two forms of post-transcriptional adenylation seen in the small non-coding RNA global sequencing experiment. Additional replicates to the sequencing of 293TREX TENT2 control and KO clone alleles (S4B) have been added, and a TENT2 immunoblot for control and TENT2 KO clones has been updated (S4C) which includes a loading control in GAPDH. Details in the text and discussion have been added to highlight the rationale for the 2 hr EU labeling experiment which produced the global sequencing dataset, and discussion around uridylation seen in datasets has been updated to reflect the unknown origin of this uridylation. The model figure has been adjusted to this end (7E).

