## Supplementary Figures and Tables for "Widespread mono- and oligoadenylation direct small noncoding RNA maturation versus degradation fates"

**SUPPLEMENTARY DATA**

### Widespread mono- and oligo-adenylation direct maturation versus degradation decisions during human small non-coding RNA biogenesis

**Cody Ocheltree, Blake Skrable, Anastasia Pimentel, Timothy Nicholson-Shaw, Suzanne R. Lee and Jens Lykke-Andersen***

**Correspondence authors:** Jens Lykke-Andersen

**This PDF file includes:**

Supporting text

Figures S1 to S6

Tables S1 to S6

#### Supplementary Figures and Tables

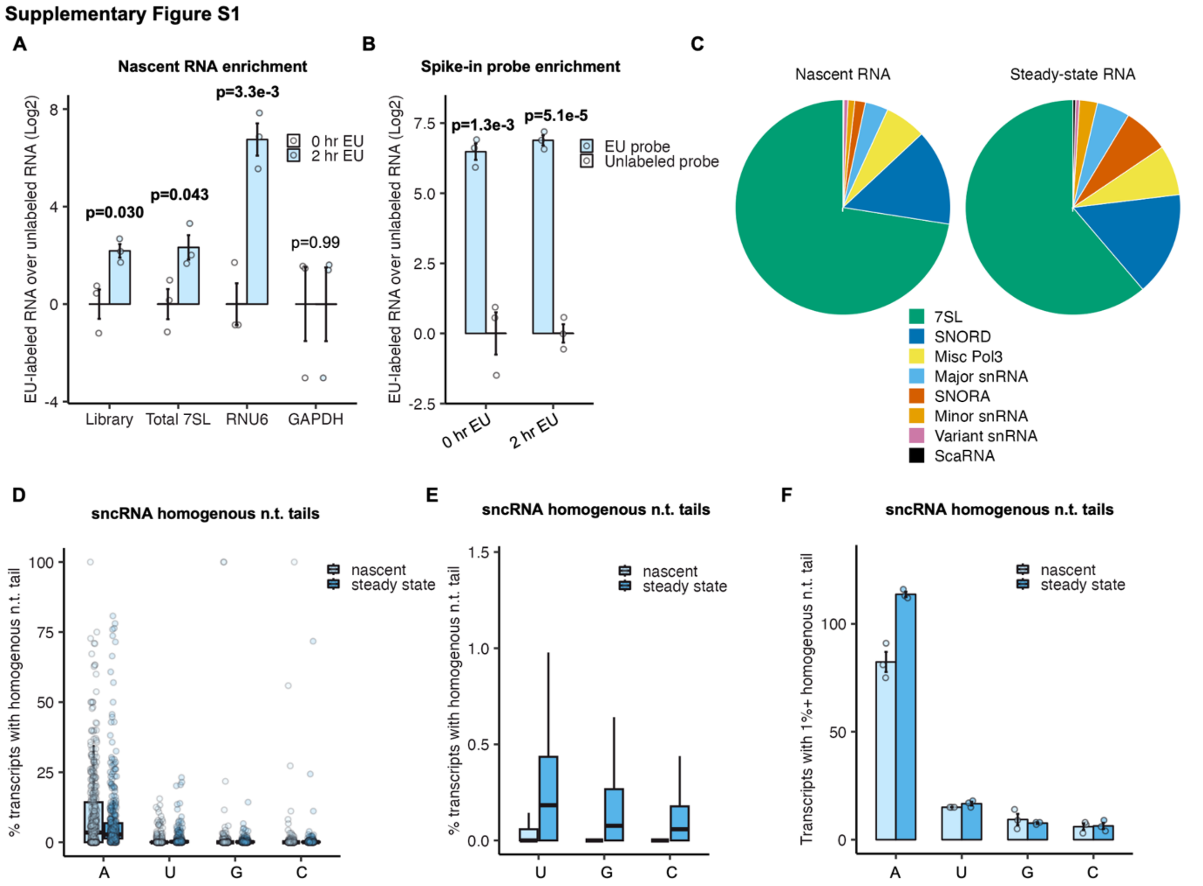


**Supplementary Figure S1: Global enrichment of nascently-transcribed sncRNAs**. **(A)** Efficiency of nascent sncRNA capture and size selection monitored by RT-qPCR for indicated RNAs from 2 hr EU- relative to DMSO (0 hr) treated samples. The library sample represents the total library monitored by RT-qPCR using primers against ligation linkers. Data is represented as mean +/- SEM and significance was determined by two-sample two-tailed t-test with p<0.05 shown in bold (n=3 per transcript). **(B)** Efficiency of nascent RNA capture monitored by RT-qPCR for EU-labeled versus unlabeled spike-in β-globin probes. Data is represented as mean +/- SEM with and significance was determined by two-sample two-tailed t-test with p<0.05 shown in bold (n=3 per transcript). **(C)** The proportion of reads aligned to each representative sncRNA class in nascent and steady state libraries. SNORA and SNORD: box H/ACA and box C/D snoRNAs, respectively. **(D)** The percent of transcripts with homogenous post-transcriptional A-, U-, G- or C-tails in nascent and steady state conditions (n=142 per condition). **(E)** Data from panel *D* plotted as ranges of percentages of transcripts with homogenous post-transcriptional U-, G- and C-tails in nascent and steady state conditions. **(F)**

Number of unique transcripts with 1% or greater homogenous post-transcriptional nucleotide tails in nascent and steady state conditions. Data is represented as mean +/- SEM (n=3 per condition).

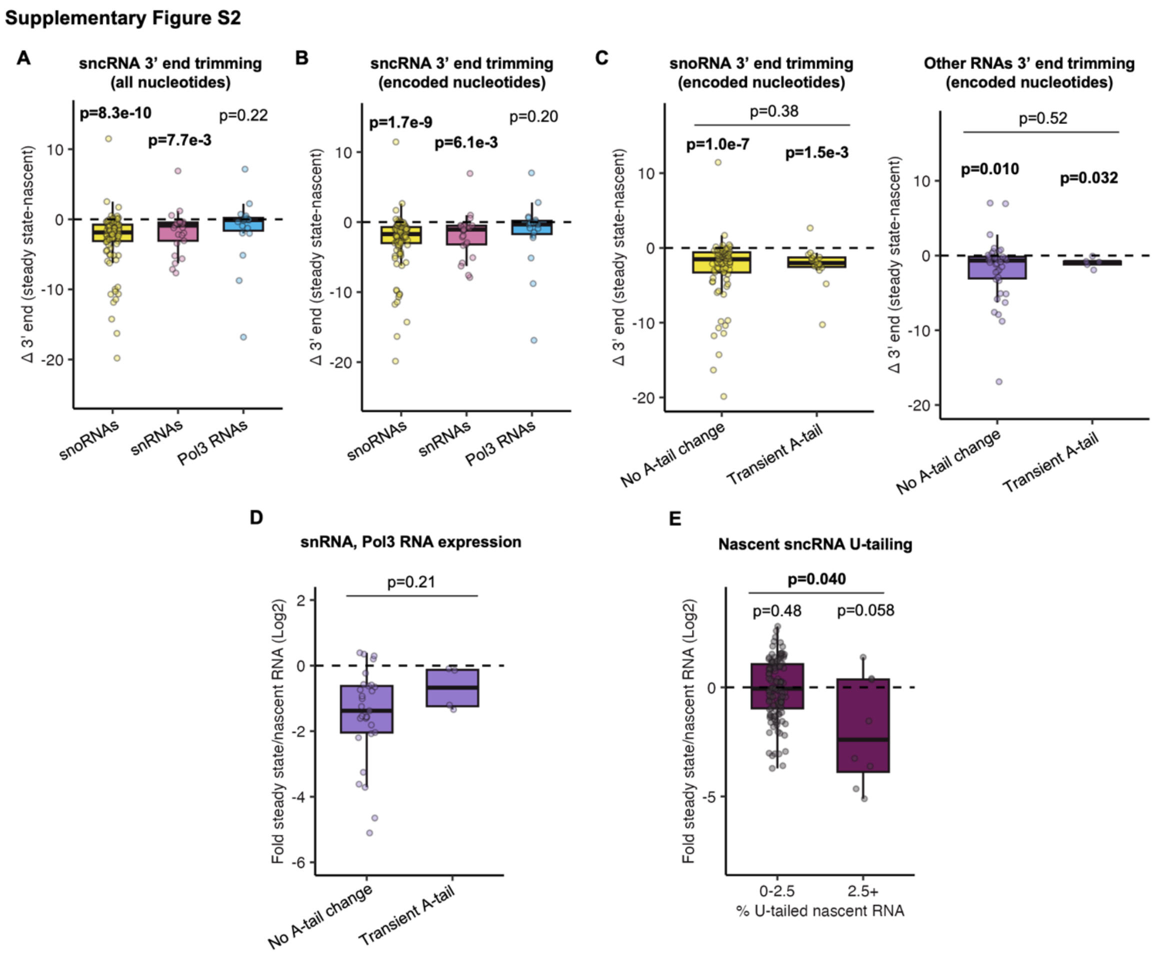


**Supplementary Figure S2: 3′ end trimming and stability of tailed sncRNAs. (A)** Plot from Figure 2A with individual transcripts visualized. **(B)** Plot from Figure 2B with individual transcripts visualized. **(C)** Box plots showing 3' end trimming of snoRNAs (left: no A-tail change n=80, transient A-tail n=18) and snRNAs/Pol-III RNAs (right: no A-tail change n=39, transient A-tail n=5) as measured by the difference in the mean 3′-end positions of transcripts in steady-state versus nascent RNA populations. RNAs that saw transient 3′ A-tailing, as measured by a significantly ($p<0.05$; two-sample two-tailed t-test) higher fraction of A-tailed molecules in nascent over steady state populations, are compared to other RNAs. Significance for each group were determined by a one-sample two-tailed t-test against mu=0 with $p<0.05$ in bold.

Significance between groups was determined by a two-sample KS test with p<0.05 in bold. **(D)** Box plots showing relative stabilities of snRNA/Pol-III RNAs as measured by log2 ratios of levels in steady-state over nascent conditions quantified using DESeq2. RNAs that saw transient 3' A-tailing, as measured by a significantly (p<0.05; two-sample two-tailed t-test) higher fraction of A-tailed molecules in nascent over steady state populations, are compared to other RNAs. Significance between groups was determined by a two-sample KS test with p<0.05 in bold (no A-tail change n=29, transient A-tail n=4). **(E)** Same as panel *D* but binning sncRNAs by % U-tailing. Significance for each group were determined by a one-sample two-tailed t-test against mu=0 with p<0.05 in bold. Significance between groups was determined by a two-sample KS test with p<0.05 in bold (0-2.5% n=121, 2.5%+ n=8).

**Supplementary Figure S3**

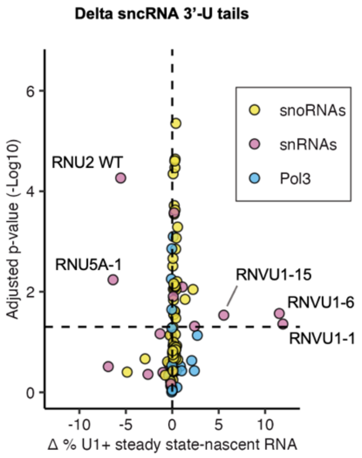


**Supplementary Figure S3: Change in nascent and steady-state sncRNA U tails.** The difference in U-tails from steady state to nascent conditions plotted against –Log10 p-values. The horizontal dashed line represents p=0.05

**Supplementary Figure S4**

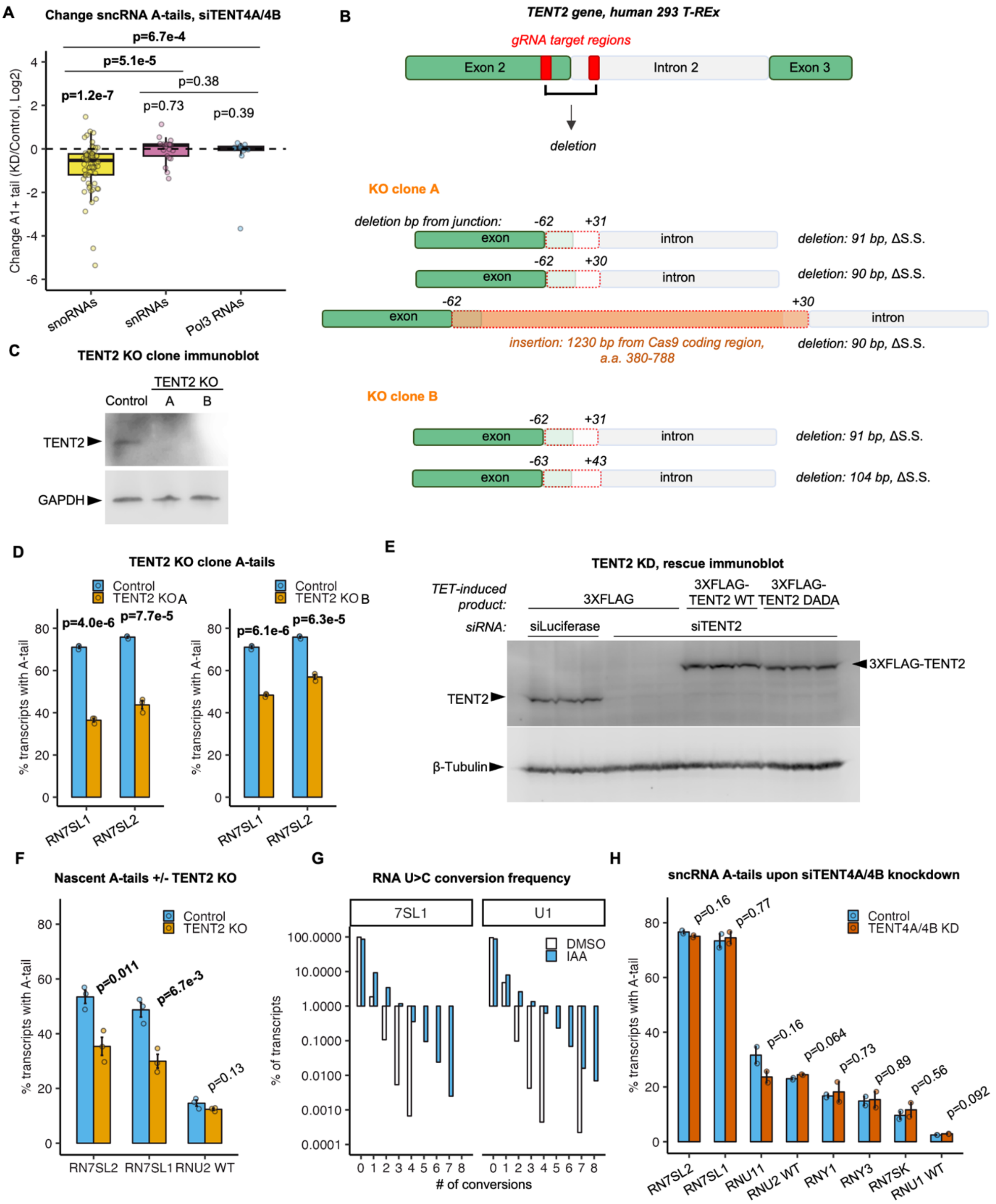


**Supplementary Figure S4: The impact of TENT2 and TENT4A/4B depletion on sncRNA A-tailing. (A)** Plot from Figure 4A with individual transcripts visualized. **(B)** Schematic of *TENT2* genomic region targeted by CRISPR-Cas9 guide (g)RNAs. *TENT2* gene alleles in KO clones A and B determined by PCR amplification of the surrounding region followed by Sanger and nanopore sequencing are illustrated; numbers above alleles refer to the position of the genomic deletion relative to the exon2-intron2 5' splice site. **(C)** Western blot for TENT2 in TENT2 KO clones A and B and a control clone that saw no TENT2 depletion. GAPDH is shown as a loading control. **(D)** Percent A-tailing of 7SL RNAs for two TENT2 KO clones. Data is represented as mean +/- SEM and significance was determined by two-sample two-tailed t-test with $p<0.05$ in bold (n=3 for each transcript). Clone A was used in all subsequent analyses. **(E)** Western blot of TENT2 protein following siRNA knockdown in 293 T-REx cells expressing exogenous 3XFLAG- or 3XFLAG-TENT2 in biological triplicate. 3XFLAG-TENT2 WT or catalytic mutant (DADA) were induced with tetracycline. b-tubulin is shown as a loading control. **(F)** Percent A-tailing of nascent 7SL and U2 RNAs from control or TENT2 KO cells metabolically labeled with s4U. Reads containing 2 or more U>C conversions after IAA-treatment were considered nascently-transcribed. Data is represented as mean +/- SEM and significance was determined by two-sample two-tailed t-test with $p<0.05$ in bold (n=3 for each transcript). **(G)** The percent of reads with 0 to 8 U>C conversions for 7SL1 and U1 RNAs after DMSO or IAA-treatment (n=1 per transcript). No bar represents none detected. **(H)** Percent A-tailing of select Pol-III/snRNAs during control or TENT4A/4B knock-down (KD) conditions (data from Lim 2018). Data is represented as mean +/- SEM and significance was determined by two-sample two-tailed t-test with $p<0.05$ in bold (n=3 for each transcript).

Supplementary Figure S5

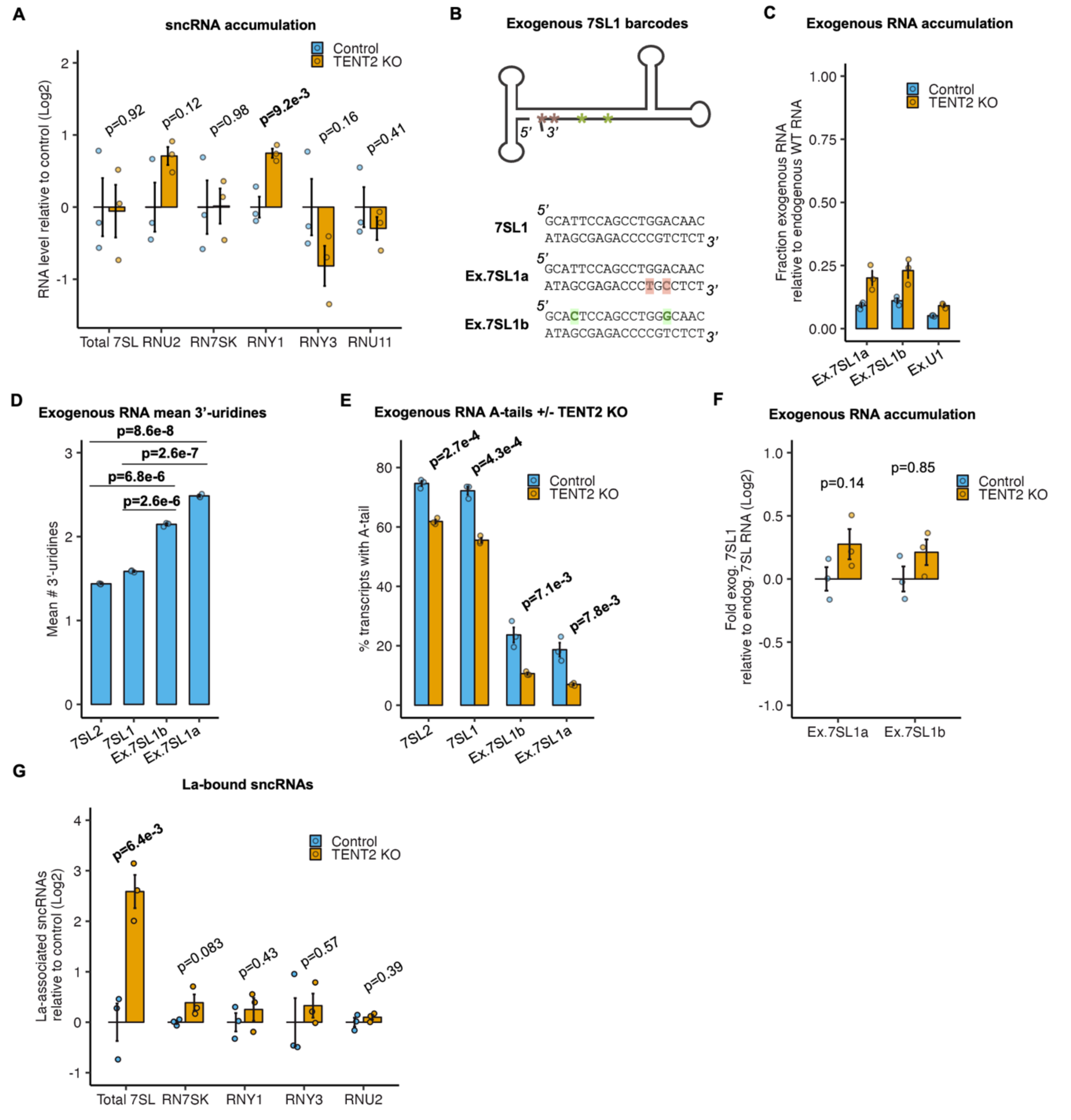


**Supplementary Figure S5: Exogenous 7SL1 RNAs accumulate with more 3'-uridines than endogenous 7SL RNAs. (A)** Impact of TENT2 KO on sncRNA levels monitored by RT-qPCR. Levels were normalized against U1 snRNA. Data is represented as mean +/- SEM and significance between groups was determined by two-sample two-tailed t-test with $p<0.05$ in bold (n=3 for each transcript). **(B)** A schematic representing the location of two point-mutations in exogenous 7SL1 RNAs *a* and *b* generated in order to distinguish exogenous from endogenous 7SL1 RNAs by sequencing. The 38 3'-terminal nucleotides of the endogenous and exogenous 7SL1 RNAs are shown below the schematic. **(C)** The abundance of exogenous 7SL1 and U1 RNAs relative to the endogenous 7SL1/2 and U1 RNAs, respectively, in control and TENT2 KO conditions monitored by gene-specific sequencing. Data is represented as mean +/- SEM (n=3 for each transcript). **(D)** Mean number of 3′-uridines of endogenous and exogenous 7SL RNAs. Data is represented as mean +/- SEM and significance between groups was determined by two-sample two-tailed t-test with $p<0.05$ in bold (n=3 for each transcript). **(E)** Percent A-tailing of endogenous and exogenous 7SL RNAs in control or TENT2 KO cells. Data is represented as mean +/- SEM and significance between groups was determined by two-sample two-tailed t-test with $p<0.05$ in bold (n=3 for each transcript). **(F)** Levels of exogenous 7SL1 RNAs relative to endogenous 7SL RNAs in control or TENT2 KO cells, normalized to the relative levels of exogenous U1 versus endogenous U1 RNAs. Data is represented as mean +/- SEM and significance between groups was determined by two-sample two-tailed t-test with $p<0.05$ in bold (n=3 for each transcript). **(G)** Levels of sncRNAs associated with La in control versus TENT2 KO conditions monitored by IP followed by RT-qPCR of sncRNAs relative to U1 snRNA and normalized against the IgG IP controls. Data is represented as mean +/- SEM and p-value was determined by a two-sample two-tailed t-test, with $p<0.05$ indicated in bold (n=3 for each transcript).

**Supplementary Figure S6**

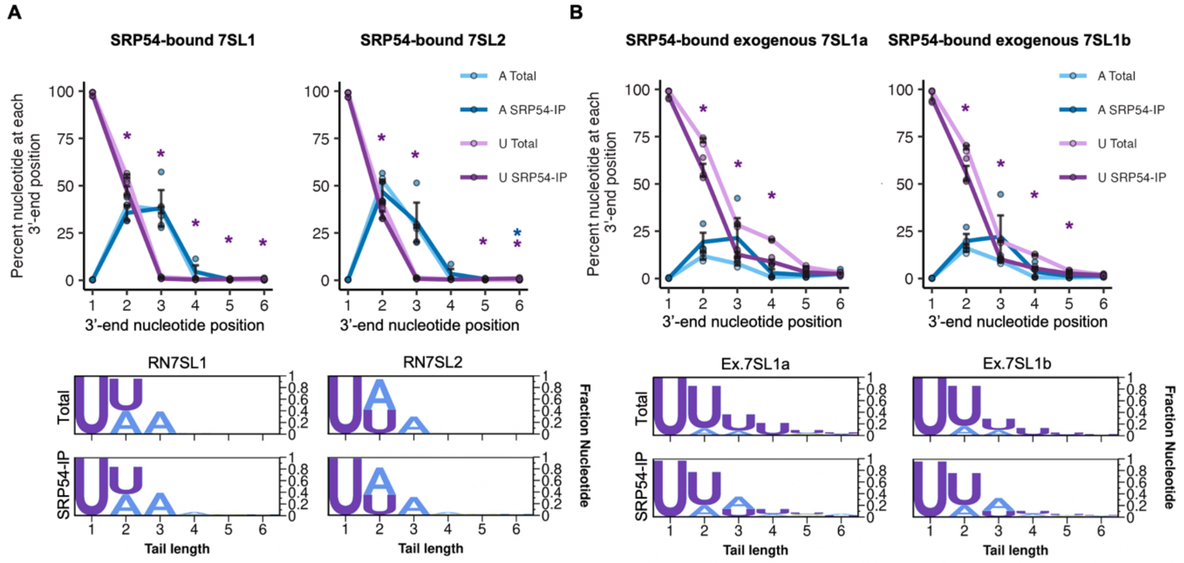


**Supplementary Figure S6: SRP54-associated 7SL RNAs are de-enriched for 3′-uridines. (A)** 3' end nucleotide compositions of steady-state versus SRP54-associated endogenous 7SL RNAs shown as line plots (top) and logo plots (bottom). Nucleotide positions are shown relative to the first nucleotide of the 7SL termination sequence, defined as position +1. Data is represented as mean +/- SEM and p-values between IP and input groups were determined by two-sample two-tailed t-tests, with $p<0.05$ indicated by asterisks (n=3 for each condition). Purple lines and asterisks represent U-tails, and blue lines/asterisks represent A-tails. **(B)** Same analysis as panel *A*, but monitoring exogenous 7SL1 RNAs.

**Supplementary Table S1.** DNA and RNA primers and sequences.

| **RNA adapter** | **sequence** |
|---|---|
| AG-10N | /5Phos/AGNNNNNNNNNNAGAUCGGAAGAGCGUCGUG/3SpC3/ |
| AG-11N | /5Phos/AGNNNNNNNNNNNAGAUCGGAAGAGCGUCGUG/3SpC3/ |
| AG-15N | /5Phos/AGNNNNNNNNNNNNNNNAGAUCGGAAGAGCGUCGUG/3SpC3/ |
| AG-16N | /5Phos/AGNNNNNNNNNNNNNNNNAGAUCGGAAGAGCGUCGUG/3SpC3/ |
| **5' cDNA adapter** | **sequence** |
| 3Tr3 | /5Phos/AGATCGGAAGAGCACACGTCTG/3SpC3/ |
| **library prep and sequencing** | **sequence** |
| AR17 | ACACGACGCTCTTCCGA |
| RC_3Tr3 | CAGACGTGTGCTCTTCCGATCT |
| RT_probe | AGGCAGAATCCAGATGC |
| TruSeq_D501 | AATGATACGGCGACCACCGAGATCTACAC TATAGCCT ACACTCTTTCCCTACACGACGCTCTTCCGATCT |
| TruSeq_D502 | AATGATACGGCGACCACCGAGATCTACAC ATAGAGGC ACACTCTTTCCCTACACGACGCTCTTCCGATCT |
| TruSeq_D503 | AATGATACGGCGACCACCGAGATCTACAC CCTATCCT ACACTCTTTCCCTACACGACGCTCTTCCGATCT |
| TruSeq_D504 | AATGATACGGCGACCACCGAGATCTACAC GGCTCTGA ACACTCTTTCCCTACACGACGCTCTTCCGATCT |
| TruSeq_D701 | CAAGCAGAAGACGGCATACGAGAT CGAGTAAT GTGACTGGAGTTCAGACGTGTGCTCTTCCGATC |
| TruSeq_D702 | CAAGCAGAAGACGGCATACGAGAT TCTCCGGA GTGACTGGAGTTCAGACGTGTGCTCTTCCGATC |
| TruSeq_D703 | CAAGCAGAAGACGGCATACGAGAT AATGAGCG GTGACTGGAGTTCAGACGTGTGCTCTTCCGATC |
| **gene specific primers for 3' end library preparation** | **sequence** |
| FU1-01 | CAGACGTGTGCTCTTCCGATCT ATCG ATGATCACGAAGGTGGTTTT |
| FU1-02 | CAGACGTGTGCTCTTCCGATCT GAACG ATGATCACGAAGGTGGTTTT |
| FU1-03 | CAGACGTGTGCTCTTCCGATCT GTTGTA ATGATCACGAAGGTGGTTTT |

| | |
|---|---|
| FU1-04 | CAGACGTGTGCTCTTCCGATCT CTACCAT ATGATCACGAAGGTGGTTTT |
| FU1-05 | CAGACGTGTGCTCTTCCGATCT TGGCTTCA ATGATCACGAAGGTGGTTTT |
| FU1-06 | CAGACGTGTGCTCTTCCGATCT ACTACGTGT ATGATCACGAAGGTGGTTTT |
| FU2-01 | CAGACGTGTGCTCTTCCGATCT TACG GGAGATGGAATAGGAGCTTGC |
| FU2-02 | CAGACGTGTGCTCTTCCGATCT CTGCT GGAGATGGAATAGGAGCTTGC |
| FU2-03 | CAGACGTGTGCTCTTCCGATCT CAATCA GGAGATGGAATAGGAGCTTGC |
| FU2-04 | CAGACGTGTGCTCTTCCGATCT GTAGATG GGAGATGGAATAGGAGCTTGC |
| FU2-05 | CAGACGTGTGCTCTTCCGATCT AGCCATAG GGAGATGGAATAGGAGCTTGC |
| FU2-06 | CAGACGTGTGCTCTTCCGATCT TGTACAGCA GGAGATGGAATAGGAGCTTGC |
| F7SL-01 | CAGACGTGTGCTCTTCCGATCT ATCG TAAGTTCGGCATCAATATGGT |
| F7SL-02 | CAGACGTGTGCTCTTCCGATCT GAACG TAAGTTCGGCATCAATATGGT |
| F7SL-03 | CAGACGTGTGCTCTTCCGATCT CGTATC TAAGTTCGGCATCAATATGGT |
| F7SL-04 | CAGACGTGTGCTCTTCCGATCT TCTGAAG TAAGTTCGGCATCAATATGGT |
| F7SL-05 | CAGACGTGTGCTCTTCCGATCT ATAGGCCA TAAGTTCGGCATCAATATGGT |
| F7SL-06 | CAGACGTGTGCTCTTCCGATCT GTCGAATCG TAAGTTCGGCATCAATATGGT |
| F7SK-01 | CAGACGTGTGCTCTTCCGATCT CTGA GCTCTCAAGGTCCATTTGTAG |
| F7SK-02 | CAGACGTGTGCTCTTCCGATCT TTACG GCTCTCAAGGTCCATTTGTAG |
| F7SK-03 | CAGACGTGTGCTCTTCCGATCT GAATAC GCTCTCAAGGTCCATTTGTAG |
| F7SK-04 | CAGACGTGTGCTCTTCCGATCT AATGCGT GCTCTCAAGGTCCATTTGTAG |
| F7SK-05 | CAGACGTGTGCTCTTCCGATCT TGCATTAA GCTCTCAAGGTCCATTTGTAG |
| F7SK-06 | CAGACGTGTGCTCTTCCGATCT ACATCAGTA GCTCTCAAGGTCCATTTGTAG |
| FY1-01 | CAGACGTGTGCTCTTCCGATCT CTGA GCTGGTCCGAAGGTAGTGAG |
| FY1-02 | CAGACGTGTGCTCTTCCGATCT TTACG GCTGGTCCGAAGGTAGTGAG |
| FY1-03 | CAGACGTGTGCTCTTCCGATCT GAATAC GCTGGTCCGAAGGTAGTGAG |
| FY1-04 | CAGACGTGTGCTCTTCCGATCT AATGCGT GCTGGTCCGAAGGTAGTGAG |

| | |
|---|---|
| FY1-05 | CAGACGTGTGCTCTTCCGATCT TGCATTAA GCTGGTCCGAAGGTAGTGAG |
| FY1-06 | CAGACGTGTGCTCTTCCGATCT ACATCAGTA GCTGGTCCGAAGGTAGTGAG |
| FY3-01 | CAGACGTGTGCTCTTCCGATCT CTGA GCTGGTCCGAGTGCAGTG |
| FY3-02 | CAGACGTGTGCTCTTCCGATCT TTACG GCTGGTCCGAGTGCAGTG |
| FY3-03 | CAGACGTGTGCTCTTCCGATCT GAATAC GCTGGTCCGAGTGCAGTG |
| FY3-04 | CAGACGTGTGCTCTTCCGATCT AATGCGT GCTGGTCCGAGTGCAGTG |
| FY3-05 | CAGACGTGTGCTCTTCCGATCT TGCATTAA GCTGGTCCGAGTGCAGTG |
| FY3-06 | CAGACGTGTGCTCTTCCGATCT ACATCAGTA GCTGGTCCGAGTGCAGTG |
| FU2-01 | CAGACGTGTGCTCTTCCGATCT TACG GGAGATGGAATAGGAGCTTGC |
| FU2-02 | CAGACGTGTGCTCTTCCGATCT CTGCT GGAGATGGAATAGGAGCTTGC |
| FU2-03 | CAGACGTGTGCTCTTCCGATCT CAATCA GGAGATGGAATAGGAGCTTGC |
| FU2-04 | CAGACGTGTGCTCTTCCGATCT GTAGATG GGAGATGGAATAGGAGCTTGC |
| FU2-05 | CAGACGTGTGCTCTTCCGATCT AGCCATAG GGAGATGGAATAGGAGCTTGC |
| FU2-06 | CAGACGTGTGCTCTTCCGATCT TGTACAGCA GGAGATGGAATAGGAGCTTGC |
| FU11-04 | CAGACGTGTGCTCTTCCGATCT ACGACTG CGACATCAAGAGATTTCGGAAGC |
| **qPCR primers** | **sequence** |
| U1_F | GCACTCCGGATGTGCTGACCC |
| U1_R | CAGGGGAAAGCGCGAACGCAG |
| 7SK_F | GAGGGCGATCTGGCTGCGACAT |
| 7SK_R | ACATGGAGCGGTGAGGGAGGAA |
| EU-labeled_Δ_β-globin_probe_F | GCAACGTGCTGGTCTGTGT |
| EU-labeled_Δ_β-globin_probe_R | CAGGGCATTAGCCACACC |
| Unlabeled_Δ_β- | GCAACCTCAAACAGACACCA |

| | |
|---|---|
| globin_probe_F | |
| Unlabeled_Δ_β-globin_probe_R | CCTCACCACCAACTTCATCC |
| 7SL_F | GGAGTTCTGGGCTGTAGTGC |
| 7SL_R | ATCAGCACGGGAGTTTTGAC |
| U6_F | GCTTCGGCAGCACATATACTAAAAT |
| U6_R | CGCTTCACGAATTTGCGTGTCAT |
| GAPDH_F | TGGCGTTGTGTGCTGAAGAT |
| GAPDH_R | TAACGGGCCAGAACATTGG |
| Y1_F | GCTGGTCCGAAGGTAGTGAG |
| Y1_R | AGTCAAGTGCAGTAGTGAGAAGG |
| Y3_F | GCTGGTCCGAGTGCAGTG |
| Y3_R | AAAAGGCTAGTCAAGTGAAGCAG |
| U2_F | CTCGGCCTTTTGGCTAAGAT |
| U2_R | ACGGAGCAAGCTCCTATTCC |
| **cloning primers** | **sequence** |
| endogenous_7SL1_F | TCGAGCTCGCCAGAATGCAGTGCTCAAAG |
| endogenous_7SL1_R | AGGATCCCCGTCTCTTGAGAGTCCAAAATTAAC |
| pUC19_F | TCTCAAGAGACGGGGATCCTCTAGAGTC |
| pUC19_R | CTGCATTCTGGCGAGCTCGAATTCCTATAG |
| 7SL1a_barcode_F | AGCGAGACCCTGCCTCTTTTGAAC |
| 7SL1a_barcode_R | ATGTTGCCCAGGCTGGAG |
| 7SL1b_barcode_F | CCTGGACAACATAGCGAGACCCCG |
| 7SL1b_barcode_R | CTGGAATGCAGTGGCTATTCACAG |
| **gRNAs** | **sequence** |
| TENT2_grna1_sense | CACCGTGAATAAACAGTAGGTGACA |
| TENT2_grna1_comp | AAACTGTCACCTACTGTTTATTCA |
| TENT2_grna2_sense | CACCGAATGAAATTTACCAACAACA |
| TENT2_grna2_comp | AAACTGTTGTTGGTAAATTTCATT |
| **genotyping** | **sequence** |
| TENT2_ex2_F | CCACCCTTCACTCCAAATCA |

| | |
|---|---|
| TENT2_ex3_R | CGAAATAATGGGGAAGCTGA |
| **siRNAs** | **sequence** |
| siControl_Luciferase | CGUACGCGGAAUACUUCGAUU |
| siTENT2 | CGUUAGUGCUGGUGAUUAAUU |
| **exogenous expression transcripts** | **sequence** |
| Barcoded U1 WT | ATACTTACCTGGCAGGGGAGATACCATGATCACGAAGGTGGTTTTCCCAGGGCGAGGCTTATCCATTGCACTCCGGATGTGCTGACCCCTGCGATTTCCCCAAAGCTGGGAAACTCGACTGCATAATTTGTGGTAGTGGGGGACTGCGTTCGCGCTTTCCCCTG |
| Barcoded 7SL1a | GCCGGGCGCGGTGGCGCGTGCCTGTAGTCCCAGCTACTCGGGAGGCTGAGGCTGGAGGATCGCTTGAGTCCAGGAGTTCTGGGCTGTAGTGCGCTATGCCGATCGGGTGTCCGCACTAAGTTCGGCATCAATATGGTGACCTCCCGGGAGCGGGGGACCACCAGGTTGCCTAAGGAGGGGTGAACCGGCCCAGGTCGGAAACGGAGCAGGTCAAAACTCCCGTGCTGATCAGTAGTGGGATCGCGCCTGTGAATAGCCACTGCACTCCAGCCTGGGCAACATAGCGAGACCCTGCCTCT |
| Barcoded 7SL1b | GCCGGGCGCGGTGGCGCGTGCCTGTAGTCCCAGCTACTCGGGAGGCTGAGGCTGGAGGATCGCTTGAGTCCAGGAGTTCTGGGCTGTAGTGCGCTATGCCGATCGGGTGTCCGCACTAAGTTCGGCATCAATATGGTGACCTCCCGGGAGCGGGGGACCACCAGGTTGCCTAAGGAGGGGTGAACCGGCCCAGGTCGGAAACGGAGCAGGTCAAAACTCCCGTGCTGATCAGTAGTGGGATCGCGCCTGTGAATAGCCACTGCATTCCAGCCTGGACAACATAGCGAGACCCCGTCTCT |
| **spike-in probes** | **sequence** |
| EU-labeled Δ β-globin probe | ATCCTGAGAACTTCAGGCTCCTGGGCAACGTGCTGGTCTGTGTGCTGGCCCATCACTTTGGCAAAGAATTCACCCCACCAGTGCAGGCTGCCTATCAGAAAGTGGTGGCTGGTGTGGCTAATGCCCTGGCCCACAAGTATCACTAAGCGGCCGCTCGCTTTCTTGCTGTCCAATTTCTATTAAAGGTTCCTTTGTTCCCTAAGTCCAACTACTAAACTGGGGGATATTATGAAGGGCCTTGAGCATCTGGATTCTGCC |
| Unlabeled Δ β-globin probe | ACATTTGCTTCTGACACAACTGTGTTCACTAGCAACCTCAAACAGACACCATGGACTACAAGGACGACGATGACAAGGTGCACCTGACTCCTGAGGAGAAGTCTGCCGTTACTGCCCTGTGGGGCAAGGTGAACGTGGATGAAGTTGGTGGTGAGGCCCTGGGCCGCTCGCTTTCTTGCTGTCCAATTTCTATTAAAGGTTCCTTTGTTCCCTAAGTCCAACTACTAAACTGGGGGATATTATGAAGGGCCTTGAGCATCTGGATTCTGCC |

**Supplementary Table S2.** TENT2 KO clone allele sanger sequencing.

| clone.sequence | sequence | |
|---|---|---|
| | | exon2_fwd exon3_rev exon mutation intron insertion (Cas9) |
| **Wild type TENT2** | CCACCCTTCACTCCAAATCATCAACAACATAATAACTTCTTTACCCTGTCACCTACTGTTTATTCACACCAGCAGCTTATAGATGCACAATTCAACTTTCAGAATGCAGAGTGAGTATGGTGATATTTTGGCCCATGTTGTTGGTAAATTTCATTTTCTATTTGACATAAACCCTTTTTGTATGAAACTTTTGAACATGGAGAATATGTCGTCACATTTTTTTTTGATACGGATGTATTTAATTTTTTTTCAGAGACTGGTTTTGTTTTTAAAGAAAAAATATGTATAAATTACTGTTCTTTTCTTTGATTTTGTTAGCTTGTCTAGAGCTGTGTCATTACAGCAGCTGACATATGGAAATGTCAGTCCAATACAGACCTCAGCTTCCCCATTATTTCGA | |
| **TENT2 Control.1** | CCACCCTTCACTCCAAATCATCAACAACATAATAACTTCTTTACCCTGTCACCTACTGTTTATTCACACCAGCAGCTTATAGATGCACAATTCAACTTTCAGAATGCAGAGTGAGTATGGTGATATTTTGGCCCATGTTGTTGGTAAATTTCATTTTCTATTTGACATAAACCCTTTTTGTATGAAACTTTTGAACATGGAGAATATGTCGTCACATTTTTTTTTGATACGGATGTATTTAATTTTTTTTCAGAGACTGGTTTTGTTTTTAAAGAAAAAATATGTATAAATTACTGTTCTTTTCTTTGATTTTGTTAGCTTGTCTAGAGCTGTGTCATTACAGCAGCTGACATATGGAAATGTCAGTCCAATACAGACCTCAGCTTCCCCATTATTTCGA | |

**TENT2 Control.2**

CCACCCTTCACTCCAAATCATCAACAACATAATAACTTCTTTACCCTGT
CACCTACTGTTTATTCACACCAGCAGCTTATAGATGCACAATTCAACT
TTCAGAATGCAGAGTGAGTATGGTGATATTTTGGCCCATGTTGTTGGT
AAATTTCATTTTCTATTTGACATAAACCCTTTTTGTATGAAACTTTTGAA
CATGGAGAATATGTCGTCACATTTTTTTTTGATACGGATGTATTTAATT
TTTTTTCAGAGACTGGTTTTGTTTTTAAAGAAAAAATATGTATAAATTAC
TGTTCTTTTCTTTGATTTTGTTAGCTTGTCTAGAGCTGTGTCATTACAG
CAGCTGACATATGGAAATGTCAGTCCAATACAGACCTCAGCTTCCCC
ATTATTTCGA

**TENT2 Control.3**

CCACCCTTCACTCCAAATCATCAACAACATAATAACTTCTTTACCCTGT
CACCTACTGTTTATTCACACCAGCAGCTTATAGATGCACAATTCAACT
TTCAGAATGCAGAGTGAGTATGGTGATATTTTGGCCCATGTTGTTGGT
AAATTTCATTTTCTATTTGACATAAACCCTTTTTGTATGAAACTTTTGAA
CATGGAGAATATGTCGTCACATTTTTTTTTGATACGGATGTATTTAATT
TTTTTTCAGAGACTGGTTTTGTTTTTAAAGAAAAAATATGTATAAATTAC
TGTTCTTTTCTTTGATTTTGTTAGCTTGTCTAGAGCTGTGTCATTACAG
CAGCTGACATATGGAAATGTCAGTCCAATACAGACCTCAGCTTCCCC
ATTATTTCGA

**TENT2 Control.4**

CCACCCTTCACTCCAAATCATCAACAACATAATAACTTCTTTACCCTGT
CACCTACTGTTTATTCACACCAGCAGCTTATAGATGCACAATTCAACT
TTCAGAATGCAGAGTGAGTATGGTGATATTTTGGCCCATGTTGTTGGT
AAATTTCATTTTCTATTTGACATAAACCCTTTTTGTATGAAACTTTTGAA
CATGGAGAATATGTCGTCACATTTTTTTTTGATACGGATGTATTTAATT
TTTTTTCAGAGACTGGTTTTGTTTTTAAAGAAAAAATATGTATAAATTAC
TGTTCTTTTCTTTGATTTTGTTAGCTTGTCTAGAGCTGTGTCATTACAG
CAGCTGACATATGGAAATGTCAGTCCAATACAGACCTCAGCTTCCCC
ATTATTTCGA

**TENT2 KOA.1**

CCACCCTTCACTCCAAATCATCAACAACATAATAACTTCTTTACCCTGT
[91 bp deletion]
TTGGTAAATTTCATTTTCTATTTGACATAAACCCTTTTTGTATGAAACTT
TTGAACATGGAGAATATGTCGTCACATTTTTTTTTGATACGGATGTATT
TAATTTTTTTTCAGAGACTGGTTTTGTTTTTAAAGAAAAAATATGTATAA
ATTACTGTTCTTTTCTTTGATTTTGTTAGCTTGTCTAGAGCTGTGTCAT
TACAGCAGCTGACATATGGAAATGTCAGTCCAATACAGACCTCAGCTT
CCCCATTATTTCGA

**TENT2 KO A.2**

TGGAAGATATCGTGCTGACCCTGACACTGTTTGAGGACAGAGAGATG
ATCGAGGAACGGCTGAAAACCTATGCCCACCTGTTCGACGACAAAGT
GATGAAGCAGCTGAAGCGGCGGAGATACACCGGCTGGGGCAGGCT
GAGCCGGAAGCTGATCAACGGCATCCGGGACAAGCAGTCCGGCAAG
ACAATCCTGGATTTCCTGAAGTCCGACGGCTTCGCCAACAGAAACTT
CATGCAGCTGATCCACGACGACAGCCTGACCTTTAAAGAGGACATCC
AGAAAGCCCAGGTGTCCGGCCAGGGCGATAGCCTGCACGAGCACAT
TGCCAATCTGGCCGGCAGCCCCGCCATTAAGAAGGGCATCCTGCAG
ACAGTGAAGGTGGTGGACGAGCTCGTGAAAGTGATGGGCCGGCACA
AGCCCGAGAACATCGTGATCGAAATGGCCAGAGAGAACCAGACCAC
CCAGAAGGGACAGAAGAACAGCCGCGAGAGAATGAAGCGGATCGAA
GAGGGCATCACTACTGTTTATTCACACCAGCAGCTTATAGATGCACAA
TTCAACTTTCAGAATGCAGAGTGAGTATGGTGATATTTTGGCCCATGT
TTGTTGGTAAATTTCATTTTCTATTTGACATAAACCCTTTTTGTATGAAA
CTTTTGAACATGGAGAATATGTCGTCACATTTTTTTTTGATACGGATGT
ATTTAATTTTTTTTCAGAGACTGGTTTTGTTTTTAAAGAAAAAATATGTA
TAAATTACTGTTCTTTTCTTTGATTTTGTTAGCTTGTCTAGAGCTGTGT
CATTACAGCAGCTGACATATGGAAATGTCAGTCCAATACAGACCTCAG
CTTCCCCATTATTTCGA

**TENT2 KO A.3**

CCACCCTTCACTCCAAATCATCAACAACATAATAACTTCTTTACCCTGT
CCTGGAAAAGATGGACGGCACCGAGGAACTGCTCGTGAAGCTGAAC
AGAGAGGACCTGCTGCGGAAGCAGCGGACCTTCGACAACGGCAGCA
TCCCCCACCAGATCCACCTGGGAGAGCTGCACGCCATTCTGCGGCG
GCAGGAAGATTTTTACCCATTCCTGAAGGACAACCGGGAAAAGATCG
AGAAGATCCTGACCTTCCGCATCCCCTACTACGTGGGCCCTCTGGCC
AGGGGAAACAGCAGATTCGCCTGGATGACCAGAAAGAGCGAGGAAA
CCATCACCCCCTGGAACTTCGAGGAAGTGGTGGACAAGGGCGCTTC
CGCCCAGAGCTTCATCGAGCGGATGACCAACTTCGATAAGAACCTGC
CCAACGAGAAGGTGCTGCCCAAGCACAGCCTGCTGTACGAGTACTTC
ACCGTGTATAACGAGCTGACCAAAGTGAAATACGTGACCGAGGGAAT
GAGAAAGCCCGCCTTCCTGAGCGGCGAGCAGAAAAAGGCCATCGTG
GACCTGCTGTTCAAGACCAACCGGAAAGTGACCGTGAAGCAGCTGAA
AGAGGACTACTTCAAGAAAATCGAGTGCTTCGACTCCGTGGAAATCT
CCGGCGTGGAAGATCGGTTCAACGCCTCCCTGGGCACATACCACGA
TCTGCTGAAAATTATCAAGGACAAGGACTTCCTGGACAATGAGGAAAA
CGAGGACATTCTGGAAGATATCGTGCTGACCCTGACACTGTTTGAGG
ACAGAGAGATGATCGAGGAACGGCTGAAAACCTATGCCCACCTGTTC
GACGACAAAGTGATGAAGCAGCTGAAGCGGCGGAGATACACCGGCT
GGGGCAGGCTGAGCCGGAAGCTGATCAACGGCATCCGGGA

**TENT2 KO A.4**

CCACCCTTCACTCCAAATCATCAACAACATAATAACTTCTTTACCCTGT
TG [90 bp deletion]
GTTGGTAAATTTCATTTTCTATTTGACATAAACCCTTTTTGTATGAAACT
TTTGAACATGGAGAATATGTCGTCACATTTTTTTTTGATACGGATGTAT
TTAATTTTTTTTCAGAGACTGGTTTTGTTTTTAAAGAAAAAATATGTATA
AATTACTGTTCTTTTCTTTGATTTTGTTAGCTTGTCTAGAGCTGTGTCA

TTACAGCAGCTGACATATGGAAATGTCAGTCCAATACAGACCTCAGCT
TCCCCATTATTTCGA

**TENT2 KO A.5**

CCACCCTTCACTCCAAATCATCAACAACATAATAACTTCTTTACCCTGT
[91 bp deletion]
TTGGTAAATTTCATTTTCTATTTGACATAAACCCTTTTTGTATGAAACTT
TTGAACATGGAGAATATGTCGTCACATTTTTTTTTGATACGGATGTATT
TAATTTTTTTTCAGAGACTGGTTTTGTTTTTAAAGAAAAAATATGTATAA
ATTACTGTTCTTTTCTTTGATTTTGTTAGCTTGTCTAGAGCTGTGTCAT
TACAGCAGCTGACATATGGAAATGTCAGTCCAATACAGACCTCAGCTT
CCCCATTATTTCGA

**TENT2 KO A.6**

CCACCCTTCACTCCAAATCATCAACAACATAATAACTTCTTTACCCTGT
CCTGGAAAAGATGGACGGCACCGAGGAACTGCTCGTGAAGCTGAAC
AGAGAGGACCTGCTGCGGAAGCAGCGGACCTTCGACAACGGCAGCA
TCCCCCACCAGATCCACCTGGGAGAGCTGCACGCCATTCTGCGGCG
GCAGGAAGATTTTTACCCATTCCTGAAGGACAACCGGGAAAAGATCG
AGAAGATCCTGACCTTCCGCATCCCCTACTACGTGGGCCCTCTGGCC
AGGGGAAACAGCAGATTCGCCTGGATGACCAGAAAGAGCGAGGAAA
CCATCACCCCCTGGAACTTCGAGGAAGTGGTGGACAAGGGCGCTTC
CGCCCAGAGCTTCATCGAGCGGATGACCAACTTCGATAAGAACCTGC
CCAACGAGAAGGTGCTGCCCAAGCACAGCCTGCTGTACGAGTACTTC
ACCGTGTATAACGAGCTGACCAAAGTGAAATACGTGACCGAGGGAAT
GAGAAAGCCCGCCTTCCTGAGCGGCGAGCAGAAAAAGGCCATCGTG
GACCTGCTGTTCAAGACCAACCGGAAAGTGACCGTGAAGCAGCTGAA
AGAGGACTACTTCAAGAAAATCGAGTGCTTCGACTCCGTGGAAATCT
CCGGCGTGGAAGATCGGTTCAACGCCTCCCTGGGCACATACCACGA
TCTGCTGAAAATTATCAAGGACAAGGACTTCCCTGGACAATGAGGAA
AACGAGGACATTCTGGAAGATATCGTGCTGACCCTGACACTGTTTGA
GGACAGAGAGATGATCGAGGAACGGCTGAAAACCTATGCCCACCTGT
TCGACGACAAAGTGATGAAGCAGCTGAAGCGGCGGAGATAC

CCACCCTTCACTCCAAATCATCAACAACATAATAACTTCTTTACCCTGT
CCTGGAAAAGATGGACGGCACCGAGGAACTGCTCGTGAAGCTGAAC
AGAGAGGACCTGCTGCGGAAGCAGCGGACCTTCGACAACGGCAGCA
TCCCCCACCAGATCCACCTGGGAGAGCTGCACGCCATTCTGCGGCG
GCAGGAAGATTTTTACCCATTCCTGAAGGACAACCGGGAAAAGATCG
AGAAGATCCTGACCTTCCGCATCCCCTACTACGTGGGCCCTCTGGCC
AGGGGAAACAGCAGATTCGCCTGGATGACCAGAAAGAGCGAGGAAA
CCATCACCCCCTGGAACTTCGAGGAAGTGGTGGACAAGGGCGCTTC
CGCCCAGAGCTTCATCGAGCGGATGACCAACTTCGATAAGAACCTGC
CCAACGAGAAGGTGCTGCCCAAGCACAGCCTGCTGTACGAGTACTTC
ACCGTGTATAACGAGCTGACCAAAGTGAAATACGTGACCGAGGGAAT
GAGAAAGCCCGCCTTCCTGAGCGGCGAGCAGAAAAAGGCCATCGTG
GACCTGCTGTTCAAGACCAACCGGAAAGTGACCGTGAAGCAGCTGAA
AGAGGACTACTTCAAGAAAATCGAGTGCTTCGACTCCGTGGAAATCT
CCGGCGTGGAAGATCGGTTCAACGCCTCCCTGGGCACATACCACGA
**TE** TCTGCTGAAAATTATCAAGGACAAGGACTTCCTGGACAATGAGGAAAA
**NT** CGAGGACATTCTGGAAGATATCGTGCTGACCCTGACACTGTTTGAGG
**2** ACAGAGAGATGATCGAGGAACGGCTGAAAACCTATGCCCACCTGTTC
**KO** GACGACAAAGTGATGAAGCAGCTGAAGCGGCGGAGATACACCGGCT
**A.7** GGGGCAGGCTGAGCCGGAAGCTGATCAACGGCATCCGGGACAAGCA
GTCCGGCAAGACAATCCTGGATTTCCTGAAGTCCGACGGCTTCGCCA
ACAGAAACTTCATGCAGCTGATCCACGACGACAGCCTGACCTTTAAA
GAGGACATCCAGAAAGCCCAGGTGTCCGGCCAGGGCGATAGCCTGC
ACGAGCACATTGCCAATCTGGCCGGCAGCCCCGCCATTAAGAAGGG
CATCCTGCAGACAGTGAAGGTGGTGGACGAGCTCGTGAAAGTGATG
GGCCGGCACAAGCCCGAGAACATCGTGATCGAAATGGCCAGAGAGA
ACCAGACCACCCAGAAGGGACAGAAGAACAGCCGCGAGAGAATGAA
GCGGATCGAAGAGGGCATCACCTACTGTTTATTCACACCAGCAGCTT
ATAGATGCACAATTCAACTTTCAGAATGCAGAGTGAGTATGGTGATAT
TTTGGCCCATGTTTGTTGGTAAATTTCATTTTCTATTTGACATAAACCC
TTTTTGTATGAAACTTTTGAACATGGAGAATATGTCGTCACATTTTTTT
TTGATACGGATGTATTTAATTTTTTTTCAGAGACTGGTTTTGTTTTTAAA
GAAAAAATATGTATAAATTACTGTTCTTTTCTTTGATTTTGTTAGCTTGT
CTAGAGCTGTGTCATTACAGCAGCTGACATATGGAAATGTCAGTCCAA
TACAGACCTCAGCTTCCCCATTATTTCG

| | |
|---|---|
| **TENT2 KOA.8** | CCACCCTTCACTCCAAATCATCAACAACATAATAACTTCTTTACCCTGT<br>CCTGGAAAAGATGGACGGCACCGAGGAACTGCTCGTGAAGCTGAAC<br>AGAGAGGACCTGCTGCGGAAGCAGCGGACCTTCGACAACGGCAGCA<br>TCCCCCACCAGATCCACCTGGGAGAGCTGCACGCCATTCTGCGGCG<br>GCAGGAAGATTTTTACCCATTCCTGAAGGACAACCGGGAAAAGATCG<br>AGAAGATCCTGACCTTCCGCATCCCCTACTACGTGGGCCCTCTGGCC<br>AGGGGAAACAGCAGATTCGCCTGGATGACCAGAAAGAGCGAGGAAA<br>CCATCACCCCCTGGAACTTCGAGGAAGTGGTGGACAAGGGCGCTTC<br>CGCCCAGAGCTTCATCGAGCGGATGACCAACTTCGATAAGAACCTGC<br>CCAACGAGAAGGTGCTGCCCAAGCACAGCCTGCTGTACGAGTACTTC<br>ACCGTGTATAACGAGCTGACCAAAGTGAAATACGTGACCGAGGGAAT<br>GAGAAAGCCCGCCTTCCTGAGCGGCGAGCAGAAAAAGGCCATCGTG<br>GACCTGCTGTTCAAGACCAACCGGAAAGTGACCGTGAAGCAGCTGAA<br>AGAGGACTACTTCAAGAAAATCGAGTGCTTCGACTCCGTGGAAATCT<br>CCGGCGTGGAAGATCGGTTCAACGCCTCCCTGGGCACATACCACGA<br>TCTGCTGAAAATTATCAAGGACAAGGACTTCCTGGACAATGAGGAAAA<br>CGAGGACATTCTGGAAGATATCGTGCTGACCCTGACACTGTTTGAGG<br>ACAGAGAGATGATCGAGGAACGGCTGAAAACCTATGCCCACCTGTTC<br>GACGACAAAGTGATGAAGCAGCTGAAGCGGCGGAGATACACCGGCT<br>GGGGCAGGCTGAGCCGGAAGCTGATCAACGGCATCCGGGACAAGCA<br>GTCCGGCAAGACAATCCTGGATTTCCTGAAGTCCGACGGCTTCGCCA<br>ACAGAAACTTCATGCAGCTGATCCACGACGACAGCCTGACCTTTAAA<br>GAGGACATCCAGAAAGCCCAGGTGTCCGGCCAGGGCGATAGCCTGC<br>ACGAGCACATTGCCAATCTGGCCGGCAGCCCCGCCATTAAGAAGGG<br>CATCCTGCAGACAGTGAAGGTGGTGGACGAGCTCGTGAAAGTGATG<br>GGCCGGCACAAGCCCGAGAACATCGTGATCGAAATGGCCAGAGAGA<br>ACCAGACCACCCAGAAGGGACAGAAGAACAGCCGCGAGAGAATGAA<br>GCGGATCGAAGAGGGCATCACCTACTGTTTATTCACACCAGCAGCTT<br>ATAGATGCACAATTCAACTTTCAGAATGCAGAGTGAGTATGGTGATAT<br>TTTGGCCCATGTTTGTTGGTAAATTTCATTTTCTATTTGACATAAACCC<br>TTTTTGTATGAAACTTTTGAACATGGAGAATATGTCGTCACATTTTTTT<br>TTGATACGGATGTATTTAATTTTTTTTCAGAGACTGGTTTTGTTTTTAAA<br>GAAAAAATATGTATAAATTACTGTTCTTTTCTTTGATTTTGTTAGCTTGT<br>CTAGAGCTGTGTCATTACAGCAGCTGACATATGGAAATGTCAGTCCAA<br>TACAGACCTCAGCTTCCCCATTATTTCG |
| **TENT2 KOA.9** | CCACCCTTCACTCCAAATCATCAACAACATAATAACTTCTTTACCCTGT<br>TG [90 bp deletion]<br>GTTGGTAAATTTCATTTTCTATTTGACATAAACCCTTTTTGTATGAAACT<br>TTTGAACATGGAGAATATGTCGTCACATTTTTTTTTGATACGGATGTAT<br>TTAATTTTTTTTCAGAGACTGGTTTTGTTTTTAAAGAAAAAATATGTATA<br>AATTACTGTTCTTTTCTTTGATTTTGTTAGCTTGTCTAGAGCTGTGTCA<br>TTACAGCAGCTGACATATGGAAATGTCAGTCCAATACAGACCTCAGCT<br>TCCCCATTATTTCGA |

**TENT2 KO A.10**

CCACCCTTCACTCCAAATCATCAACAACATAATAACTTCTTTACCCTGT
[91 bp deletion]
TTGGTAAATTTCATTTTCTATTTGACATAAACCCTTTTTGTATGAAACTT
TTGAACATGGAGAATATGTCGTCACATTTTTTTTTGATACGGATGTATT
TAATTTTTTTTCAGAGACTGGTTTTGTTTTTAAAGAAAAAATATGTATAA
ATTACTGTTCTTTTCTTTGATTTTGTTAGCTTGTCTAGAGCTGTGTCAT
TACAGCAGCTGACATATGGAAATGTCAGTCCAATACAGACCTCAGCTT
CCCCATTATTTCGA

**TENT2 KO B.1**

CCACCCTTCACTCCAAATCATCAACAACATAATAACTTCTTTACCCTGT
[91 bp deletion]
TTGGTAAATTTCATTTTCTATTTGACATAAACCCTTTTTGTATGAAACTT
TTGAACATGGAGAATATGTCGTCACATTTTTTTTTGATACGGATGTATT
TAATTTTTTTTCAGAGACTGGTTTTGTTTTTAAAGAAAAAATATGTATAA
ATTACTGTTCTTTTCTTTGATTTTGTTAGCTTGTCTAGAGCTGTGTCAT
TACAGCAGCTGACATATGGAAATGTCAGTCCAATACAGACCTCAGCTT
CCCCATTATTTCGA

**TENT2 KO B.2**

CCACCCTTCACTCCAAATCATCAACAACATAATAACTTCTTTACCCTG
[104 bp deletion]
ATTTTCTATTTGACATAAACCCTTTTTGTATGAAACTTTTGAACATGGA
GAATATGTCGTCACATTTTTTTTTGATACGGATGTATTTAATTTTTTTTC
AGAGACTGGTTTTGTTTTTAAAGAAAAAATATGTATAAATTACTGTTCT
TTTCTTTGATTTTGTTAGCTTGTCTAGAGCTGTGTCATTACAGCAGCTG
ACATATGGAAATGTCAGTCCAATACAGACCTCAGCTTCCCCATTATTT
CGA

**TENT2 KO B.3**

CCACCCTTCACTCCAAATCATCAACAACATAATAACTTCTTTACCCTGT
[91 bp deletion]
TTGGTAAATTTCATTTTCTATTTGACATAAACCCTTTTTGTATGAAACTT
TTGAACATGGAGAATATGTCGTCACATTTTTTTTTGATACGGATGTATT
TAATTTTTTTTCAGAGACTGGTTTTGTTTTTAAAGAAAAAATATGTATAA
ATTACTGTTCTTTTCTTTGATTTTGTTAGCTTGTCTAGAGCTGTGTCAT
TACAGCAGCTGACATATGGAAATGTCAGTCCAATACAGACCTCAGCTT
CCCCATTATTTCGA

**TENT2 KO B.4**

CCACCCTTCACTCCAAATCATCAACAACATAATAACTTCTTTACCCTG
[104 bp deletion]
ATTTTCTATTTGACATAAACCCTTTTTGTATGAAACTTTTGAACATGGA
GAATATGTCGTCACATTTTTTTTTGATACGGATGTATTTAATTTTTTTTC
AGAGACTGGTTTTGTTTTTAAAGAAAAAATATGTATAAATTACTGTTCT
TTTCTTTGATTTTGTTAGCTTGTCTAGAGCTGTGTCATTACAGCAGCTG
ACATATGGAAATGTCAGTCCAATACAGACCTCAGCTTCCCCATTATTT
CGA

| | |
|---|---|
| TENT2 KO B.5 | CCACCCTTCACTCCAAATCATCAACAACATAATAACTTCTTTACCCTG<br>[104 bp deletion]<br>ATTTTCTATTTGACATAAACCCTTTTTGTATGAAACTTTTGAACATGGA<br>GAATATGTCGTCACATTTTTTTTTGATACGGATGTATTTAATTTTTTTTC<br>AGAGACTGGTTTTGTTTTTAAAGAAAAAATATGTATAAATTACTGTTCT<br>TTTCTTTGATTTTGTTAGCTTGTCTAGAGCTGTGTCATTACAGCAGCTG<br>ACATATGGAAATGTCAGTCCAATACAGACCTCAGCTTCCCCATTATTT<br>CGA |
| TENT2 KO B.6 | CCACCCTTCACTCCAAATCATCAACAACATAATAACTTCTTTACCCTG<br>[104 bp deletion]<br>ATTTTCTATTTGACATAAACCCTTTTTGTATGAAACTTTTGAACATGGA<br>GAATATGTCGTCACATTTTTTTTTGATACGGATGTATTTAATTTTTTTTC<br>AGAGACTGGTTTTGTTTTTAAAGAAAAAATATGTATAAATTACTGTTCT<br>TTTCTTTGATTTTGTTAGCTTGTCTAGAGCTGTGTCATTACAGCAGCTG<br>ACATATGGAAATGTCAGTCCAATACAGACCTCAGCTTCCCCATTATTT<br>CGA |

**Supplementary Table S3.** Change in sncRNA tailing steadystate/nascent.

| genename | steadystate/nascent % tails (log2) | pvalue | tail | type |
|---|---|---|---|---|
| SCARNA4 | -4.8663574 | 0.05538213 | A tail | snoRNAs |
| RNU5D-1 | -4.0173365 | 0.4014856 | A tail | snRNAs |
| SNORD32A | -3.8462384 | 0.34372874 | A tail | snoRNAs |
| SNORA38 | -3.5664517 | 0.41178672 | A tail | snoRNAs |
| RNU4-1 | -3.3150006 | 0.12274862 | A tail | snRNAs |
| RNU4-2 | -3.2271992 | 0.02396384 | A tail | snRNAs |
| SNORA80C | -3.0217168 | 0.43015367 | A tail | snoRNAs |
| SNORD95 | -2.9090169 | 0.30513041 | A tail | snoRNAs |
| SNORA3B | -2.8742785 | 0.43679399 | A tail | snoRNAs |
| SNORA38B | -2.7776884 | 0.28576845 | A tail | snoRNAs |
| RNU12 | -2.6438281 | 0.01969178 | A tail | snRNAs |
| SNORD17 | -2.577037 | 0.00552904 | A tail | snoRNAs |
| SNORA77B | -2.5275739 | 0.00414724 | A tail | snoRNAs |
| SNORA9 | -2.5026634 | 0.06155238 | A tail | snoRNAs |
| SNORA68 | -2.3551204 | 0.00403981 | A tail | snoRNAs |
| SNORA64 | -2.3424248 | 0.00629052 | A tail | snoRNAs |
| SNORA50C | -2.3204513 | 3.14E-05 | A tail | snoRNAs |
| SNORA23 | -2.2883535 | 0.28601952 | A tail | snoRNAs |
| SNORA4 | -2.2801569 | 0.10214294 | A tail | snoRNAs |
| SNORA74B | -2.271044 | 0.47228963 | A tail | snoRNAs |
| SNORA71A | -2.2455435 | 0.00063566 | A tail | snoRNAs |
| SNORD94 | -2.2261118 | 0.18820895 | A tail | snoRNAs |
| SNORA63 | -2.1951742 | 0.00277778 | A tail | snoRNAs |
| SNORA57 | -2.1816808 | 0.00064164 | A tail | snoRNAs |

| | | | | |
|---|---|---|---|---|
| SNORA77 | -2.1635198 | 0.02136416 | A tail | snoRNAs |
| SNORA50B | -2.0046049 | 0.40053843 | A tail | snoRNAs |
| SNORA61 | -1.8768669 | 0.28558503 | A tail | snoRNAs |
| SNORD104 | -1.8717024 | 0.01518487 | A tail | snoRNAs |
| SNORD68 | -1.7322393 | 0.01578252 | A tail | snoRNAs |
| SNORA48 | -1.7031534 | 0.01321642 | A tail | snoRNAs |
| SNORA37 | -1.6168216 | 0.53770681 | A tail | snoRNAs |
| TERC | -1.5901659 | 0.37702581 | A tail | snRNAs |
| RNU2-2P | -1.587655 | 0.1279127 | A tail | snRNAs |
| RNVU1-14 | -1.553615 | 0.30839616 | A tail | snRNAs |
| SNORD23 | -1.5314065 | 0.10387489 | A tail | snoRNAs |
| SCARNA11 | -1.4723212 | 0.55828691 | A tail | snoRNAs |
| SNORA80B | -1.4354449 | 0.22329044 | A tail | snoRNAs |
| SCARNA8 | -1.4131698 | 0.38943572 | A tail | snoRNAs |
| RMRP | -1.3996772 | 0.00043496 | A tail | Pol3 RNAs |
| SNORA52 | -1.3962248 | 0.01638198 | A tail | snoRNAs |
| SNORA71D | -1.3558244 | 0.42520216 | A tail | snoRNAs |
| SNORD13 | -1.3194127 | 0.0010123 | A tail | snoRNAs |
| SNORA80A | -1.2792311 | 0.18202696 | A tail | snoRNAs |
| SNORA10 | -1.1639536 | 0.26705944 | A tail | snoRNAs |
| SNORA47 | -1.1598987 | 0.32016636 | A tail | snoRNAs |
| SNORA73B | -1.037559 | 0.08398074 | A tail | snoRNAs |
| SNORA21 | -1.003254 | 0.40880363 | A tail | snoRNAs |
| SNORA7A | -0.9799245 | 0.00177292 | A tail | snoRNAs |
| SNORA17B | -0.9711533 | 0.05575056 | A tail | snoRNAs |
| SNORA5C | -0.9449912 | 7.98E-05 | A tail | snoRNAs |
| SNORA3A | -0.9382697 | 0.65764311 | A tail | snoRNAs |
| SNORA73A | -0.9276245 | 0.13121638 | A tail | snoRNAs |
| RNU1 WT | -0.9269585 | 0.00831115 | A tail | snRNAs |
| SNORA44 | -0.8948625 | 0.50144224 | A tail | snoRNAs |
| SNORD105B | -0.8437564 | 0.54754433 | A tail | snoRNAs |
| RNA5S | -0.8102424 | 0.03925901 | A tail | Pol3 RNAs |
| SCARNA2 | -0.8031877 | 0.6946694 | A tail | snRNAs |
| SNORD83B | -0.7588733 | 0.27172677 | A tail | snoRNAs |
| SNORA7B | -0.7289514 | 0.10719144 | A tail | snoRNAs |
| SNORA78 | -0.7150128 | 0.02063054 | A tail | snoRNAs |
| SNORA71C | -0.6926422 | 0.6526196 | A tail | snoRNAs |
| SNORD37 | -0.6677369 | 0.61912218 | A tail | snoRNAs |
| VTRNA1-2 | -0.6136795 | 0.75813914 | A tail | Pol3 RNAs |
| SCARNA12 | -0.5566653 | 0.02833995 | A tail | snoRNAs |
| SNORD55 | -0.5309672 | 0.77357621 | A tail | snoRNAs |
| AL138963.2 | -0.4756604 | 0.60863585 | A tail | snoRNAs |
| SNORA84 | -0.4600264 | 0.54203635 | A tail | snoRNAs |

| | | | | |
|---|---|---|---|---|
| SNORA16A | -0.4413719 | 0.80669794 | A tail | snoRNAs |
| AC090227.1 | -0.3609811 | 0.83746382 | A tail | snoRNAs |
| RNU6 WT | -0.3241942 | 0.13630561 | A tail | Pol3 RNAs |
| SNORD10 | -0.2483599 | 0.33954587 | A tail | snoRNAs |
| RNU5B-1 | -0.2429427 | 0.65806128 | A tail | snRNAs |
| SCARNA22 | -0.2406299 | 0.36231212 | A tail | snoRNAs |
| SNORA18 | -0.2405809 | 0.88580844 | A tail | snoRNAs |
| SNORD118 | -0.2070089 | 0.90059072 | A tail | snRNAs |
| RNVU1-27 | -0.0822484 | 0.66390803 | A tail | snRNAs |
| RNU6ATAC | -0.0115713 | 0.97232221 | A tail | Pol3 RNAs |
| SNORA80D | 0.08032266 | 0.92067362 | A tail | snoRNAs |
| SNORD3 | 0.1021973 | 0.73670069 | A tail | snRNAs |
| SNORD88A | 0.12257119 | 0.76119203 | A tail | snoRNAs |
| SNORA80E | 0.15132536 | 0.43563101 | A tail | snoRNAs |
| RN7SK | 0.19005978 | 0.18594246 | A tail | Pol3 RNAs |
| RN7SL2 | 0.22887289 | 0.02337744 | A tail | Pol3 RNAs |
| SCARNA20 | 0.23414233 | 0.47737529 | A tail | snoRNAs |
| RN7SL1 | 0.33331307 | 0.01148813 | A tail | Pol3 RNAs |
| RNA5SP19 | 0.37439552 | 0.78242611 | A tail | Pol3 RNAs |
| SNORA74D | 0.39849479 | 0.29075129 | A tail | snoRNAs |
| VTRNA1-3 | 0.40503143 | 0.63339083 | A tail | Pol3 RNAs |
| VTRNA1-1 | 0.43373573 | 0.33002179 | A tail | Pol3 RNAs |
| RN7SL3 | 0.43396773 | 0.0069469 | A tail | Pol3 RNAs |
| RNVU1-15 | 0.43922458 | 0.28389145 | A tail | snRNAs |
| RPPH1 | 0.50164145 | 0.55941255 | A tail | Pol3 RNAs |
| SNORA70 | 0.50453515 | 0.22996504 | A tail | snoRNAs |
| RNVU1-7 | 0.53016498 | 0.45454314 | A tail | snRNAs |
| SNORA67 | 0.58730907 | 0.64196184 | A tail | snoRNAs |
| SNORA51 | 0.61808295 | 0.35918728 | A tail | snoRNAs |
| RNU11 | 0.64462447 | 0.20503137 | A tail | snRNAs |
| RNVU1-6 | 0.65662053 | 0.40665407 | A tail | snRNAs |
| SNORD22 | 0.6592877 | 0.59547751 | A tail | snoRNAs |
| RNY1 | 0.82381963 | 0.0101833 | A tail | Pol3 RNAs |
| SNORA30 | 0.86723253 | 0.48225727 | A tail | snoRNAs |
| RNY3 | 0.87602758 | 0.24186091 | A tail | Pol3 RNAs |
| SCARNA10 | 0.98795733 | 0.38549951 | A tail | snoRNAs |
| SNORD88C | 1.1444289 | 0.07735554 | A tail | snoRNAs |
| SNORA65 | 1.28983492 | 0.22335363 | A tail | snoRNAs |
| RNU5A-1 | 1.42922496 | 0.17276792 | A tail | snRNAs |
| SNORA21B | 1.67705458 | 0.21046935 | A tail | snoRNAs |
| RNVU1-31 | 2.06183156 | 0.00432659 | A tail | snRNAs |
| RNU5F-1 | 2.2511881 | 0.03692243 | A tail | snRNAs |
| RNU2 WT | 2.6458099 | 0.00175266 | A tail | snRNAs |

| | | | | |
|---|---|---|---|---|
| SNORA51 | -4.2543186 | 0.39704017 | U tail | snoRNAs |
| RNU5A-1 | -3.9537064 | 0.00577042 | U tail | snRNAs |
| SNORA71D | -3.1452719 | 0.21612938 | U tail | snoRNAs |
| RNU2-2P | -2.5913807 | 0.06860731 | U tail | snRNAs |
| SCARNA20 | -2.5008827 | 0.45683529 | U tail | snoRNAs |
| RNU2 WT | -2.0650713 | 5.45E-05 | U tail | snRNAs |
| RNU5B-1 | -1.8045106 | 0.40457214 | U tail | snRNAs |
| SNORD88C | -1.6734856 | 0.24679865 | U tail | snoRNAs |
| RNVU1-14 | -1.5241806 | 0.30872349 | U tail | snRNAs |
| SNORA70 | -1.2070065 | 0.60336388 | U tail | snoRNAs |
| RNU6 WT | -1.1078762 | 0.47052276 | U tail | Pol3 RNAs |
| RNU4-1 | -1.0859542 | 0.62510272 | U tail | snRNAs |
| SNORD10 | -1.0856403 | 0.43634339 | U tail | snoRNAs |
| RNU12 | -0.9614103 | 0.07042393 | U tail | snRNAs |
| SNORA68 | -0.9490841 | 0.4416177 | U tail | snoRNAs |
| SNORA78 | -0.9154822 | 0.25912626 | U tail | snoRNAs |
| RNU5F-1 | -0.8563659 | 0.68007542 | U tail | snRNAs |
| SNORA17B | -0.8558407 | 0.42332807 | U tail | snoRNAs |
| RNVU1-27 | -0.8285383 | 0.43986971 | U tail | snRNAs |
| RNVU1-7 | -0.5795643 | 0.26994048 | U tail | snRNAs |
| SNORA52 | -0.4865257 | 0.75152018 | U tail | snoRNAs |
| SNORD88A | -0.2069668 | 0.84496067 | U tail | snoRNAs |
| SNORA50C | -0.145508 | 0.73629834 | U tail | snoRNAs |
| SNORD83B | -0.1179995 | 0.89783522 | U tail | snoRNAs |
| RNU1 WT | -0.1059562 | 0.49581552 | U tail | snRNAs |
| SNORA73B | -0.1037906 | 0.88637142 | U tail | snoRNAs |
| SCARNA12 | -0.0776996 | 0.94751851 | U tail | snoRNAs |
| RN7SK | -0.0693074 | 0.91692832 | U tail | Pol3 RNAs |
| VTRNA1-3 | -0.011594 | 0.98773195 | U tail | Pol3 RNAs |
| VTRNA1-2 | 0.18874895 | 0.79985556 | U tail | Pol3 RNAs |
| RNU4-2 | 0.22415242 | 0.87705477 | U tail | snRNAs |
| VTRNA1-1 | 0.69823758 | 0.23463219 | U tail | Pol3 RNAs |
| SNORA64 | 0.70575916 | 0.56743869 | U tail | snoRNAs |
| RNU6ATAC | 0.88549327 | 0.13412711 | U tail | Pol3 RNAs |
| RNY1 | 1.18789501 | 0.30367679 | U tail | Pol3 RNAs |
| RNVU1-15 | 1.2324027 | 0.02927157 | U tail | snRNAs |
| RNY3 | 1.82404489 | 0.07375107 | U tail | Pol3 RNAs |
| RNVU1-6 | 1.98196759 | 0.02709502 | U tail | snRNAs |
| RNVU1-31 | 2.09656502 | 0.00802485 | U tail | snRNAs |

**Supplementary Table S4.** Transient adenylation of snoRNA species.

| genename | delta %A1+ tail (nascent-steady state) | pvalue (-log10) |
|---|---|---|
| SCARNA21B | -10.219517 | 6.127261 |
| SNORD66 | -2.555663 | 4.363512 |
| SNORD33 | -2.739262 | 4.191114 |
| SNORD46 | -6.306531 | 3.920009 |
| SNORD86 | -3.942887 | 3.534375 |
| SNORD83A | -2.447758 | 3.342384 |
| SNORA5A | -2.572028 | 2.91863 |
| SNORA2C | -1.499559 | 2.806256 |
| SNORA14B | -1.468599 | 2.706313 |
| SNORD67 | -1.48834 | 2.499182 |
| SCARNA5 | -3.705964 | 2.118934 |
| SNORD34 | -2.09566 | 1.792551 |
| SNORA71E | -1.029455 | 1.634242 |
| SNORA33 | -2.412369 | 1.510838 |

**Supplementary Table S5**. Percent of sncRNA transcripts with A/U tails in the steady state.

| genename | tail | condition | % of transcripts | sem | type |
|---|---|---|---|---|---|
| RN7SL2 | A tail | steady state | 77.1791014 | 1.4718139 | Pol3 RNAs |
| RN7SL1 | A tail | steady state | 72.8881683 | 1.4429341 | Pol3 RNAs |
| RN7SL3 | A tail | steady state | 60.1205574 | 2.2024955 | Pol3 RNAs |
| SCARNA22 | A tail | steady state | 43.0102619 | 0.6183435 | snoRNAs |
| RNVU1-1 | A tail | steady state | 32.020202 | 7.4110031 | snRNAs |
| RN7SK | A tail | steady state | 30.6537378 | 0.8418933 | Pol3 RNAs |
| RNU2 WT | A tail | steady state | 30.6448257 | 3.4043259 | snRNAs |
| SCARNA20 | A tail | steady state | 27.4002833 | 0.9495136 | snoRNAs |
| RNU11 | A tail | steady state | 26.3231065 | 0.4421652 | snRNAs |
| SNORA30 | A tail | steady state | 24.3221377 | 4.889923 | snoRNAs |
| SNORA80E | A tail | steady state | 23.1120382 | 0.7935896 | snoRNAs |
| RNY1 | A tail | steady state | 21.3167916 | 1.3645324 | Pol3 RNAs |
| RNY3 | A tail | steady state | 21.27511 | 0.9996264 | Pol3 RNAs |
| SNORA80D | A tail | steady state | 20.7944847 | 0.6513961 | snoRNAs |
| RNA5SP19 | A tail | steady state | 17.2839506 | 1.0706388 | Pol3 RNAs |
| SNORD105B | A tail | steady state | 17.079395 | 3.5758475 | snoRNAs |
| SNORD36B | A tail | steady state | 15 | 7.6376262 | snoRNAs |
| RNVU1-15 | A tail | steady state | 14.1382427 | 2.871379 | snRNAs |
| SNORA7B | A tail | steady state | 13.7740407 | 0.6552278 | snoRNAs |
| SNORA67 | A tail | steady state | 12.5203481 | 0.2965132 | snoRNAs |
| SNORA5C | A tail | steady state | 12.028549 | 0.2619078 | snoRNAs |
| AL138963.2 | A tail | steady state | 11.7002538 | 1.249733 | snoRNAs |
| SNORA7A | A tail | steady state | 11.0587982 | 0.5866675 | snoRNAs |
| AC090227.1 | A tail | steady state | 10.3817983 | 2.0181275 | snoRNAs |
| SCARNA21B | A tail | steady state | 10.2195172 | 0.1919966 | snoRNAs |
| SNORA74D | A tail | steady state | 9.9007018 | 1.2329873 | snoRNAs |
| SNORD104 | A tail | steady state | 9.5905399 | 1.4321551 | snoRNAs |
| SNORA9 | A tail | steady state | 9.1166163 | 1.8119025 | snoRNAs |
| RNU6ATAC | A tail | steady state | 8.8185629 | 0.3211878 | Pol3 RNAs |
| VTRNA1-1 | A tail | steady state | 8.2284977 | 0.8775977 | Pol3 RNAs |
| SNORD88C | A tail | steady state | 7.3918985 | 0.1596619 | snoRNAs |
| SNORA57 | A tail | steady state | 7.3478174 | 0.7298511 | snoRNAs |
| SNORA80A | A tail | steady state | 7.3442056 | 0.3530451 | snoRNAs |
| RNVU1-27 | A tail | steady state | 6.5030209 | 0.2549033 | snRNAs |
| SNORD46 | A tail | steady state | 6.3065309 | 0.4251201 | snoRNAs |
| SNORA47 | A tail | steady state | 6.1889888 | 0.4380341 | snoRNAs |
| SNORA84 | A tail | steady state | 5.9225646 | 0.5338577 | snoRNAs |
| RNA5SP452 | A tail | steady state | 5.91133 | 4.3038891 | Pol3 RNAs |
| SNORA48 | A tail | steady state | 5.7516663 | 0.0722914 | snoRNAs |
| SNORA77 | A tail | steady state | 5.7146877 | 0.249973 | snoRNAs |
| SNORA61 | A tail | steady state | 5.7047962 | 0.0842388 | snoRNAs |

| | | | | | |
|---|---|---|---|---|---|
| TERC | A tail | steady state | 5.5405876 | 0.2443989 | snRNAs |
| SNORA63 | A tail | steady state | 5.497299 | 0.6668318 | snoRNAs |
| SNORD68 | A tail | steady state | 5.452255 | 1.5633981 | snoRNAs |
| SCARNA12 | A tail | steady state | 5.3473259 | 0.089979 | snoRNAs |
| SNORA4 | A tail | steady state | 5.1197596 | 0.9328874 | snoRNAs |
| SNORD88A | A tail | steady state | 5.0265988 | 0.0931053 | snoRNAs |
| RNA5SP304 | A tail | steady state | 4.9910873 | 2.6618867 | Pol3 RNAs |
| SNORD23 | A tail | steady state | 4.9384565 | 0.4051245 | snoRNAs |
| VTRNA1-3 | A tail | steady state | 4.9140139 | 0.6099629 | Pol3 RNAs |
| RNVU1-6 | A tail | steady state | 4.8918523 | 1.6887851 | snRNAs |
| SNORA21 | A tail | steady state | 4.5556208 | 0.2345026 | snoRNAs |
| SNORD83B | A tail | steady state | 4.4604838 | 0.0217324 | snoRNAs |
| SNORA78 | A tail | steady state | 4.4205977 | 0.2298836 | snoRNAs |
| SNORA44 | A tail | steady state | 4.3734759 | 0.183717 | snoRNAs |
| SCARNA2 | A tail | steady state | 4.1527646 | 1.1011697 | snRNAs |
| SNORD38A | A tail | steady state | 4.1369298 | 2.015675 | snoRNAs |
| SNORD86 | A tail | steady state | 3.9428865 | 0.3332608 | snoRNAs |
| SNORA62 | A tail | steady state | 3.7970336 | 1.7876695 | snoRNAs |
| SCARNA5 | A tail | steady state | 3.7059638 | 0.7443658 | snoRNAs |
| SNORD24 | A tail | steady state | 3.7037037 | 3.7037037 | snoRNAs |
| SCARNA10 | A tail | steady state | 3.6729162 | 0.2665881 | snoRNAs |
| SNORA18 | A tail | steady state | 3.5266851 | 0.36894 | snoRNAs |
| RMRP | A tail | steady state | 3.5232493 | 0.1308549 | Pol3 RNAs |
| SCARNA8 | A tail | steady state | 3.3972523 | 0.2885692 | snoRNAs |
| SNORA64 | A tail | steady state | 3.3816421 | 0.1366099 | snoRNAs |
| SNORA80B | A tail | steady state | 3.3506633 | 0.2010376 | snoRNAs |
| SNORD94 | A tail | steady state | 3.315943 | 0.4256595 | snoRNAs |
| SNORD3 | A tail | steady state | 3.1712174 | 0.4390445 | snRNAs |
| SNORA38B | A tail | steady state | 3.1248249 | 0.1135661 | snoRNAs |
| RNU6 WT | A tail | steady state | 3.063119 | 0.3777772 | Pol3 RNAs |
| SNORA77B | A tail | steady state | 2.9816475 | 0.0502869 | snoRNAs |
| SNORA51 | A tail | steady state | 2.9451316 | 0.2065064 | snoRNAs |
| RPPH1 | A tail | steady state | 2.8819756 | 0.0760027 | Pol3 RNAs |
| SNORA38 | A tail | steady state | 2.8136482 | 0.7274626 | snoRNAs |
| SNORD22 | A tail | steady state | 2.800182 | 0.1904219 | snoRNAs |
| SNORD33 | A tail | steady state | 2.7392618 | 0.1576775 | snoRNAs |
| SNORA65 | A tail | steady state | 2.7166675 | 0.0949939 | snoRNAs |
| SNORA17B | A tail | steady state | 2.5862798 | 0.1610027 | snoRNAs |
| SNORA5A | A tail | steady state | 2.5720278 | 0.3137285 | snoRNAs |
| SNORD66 | A tail | steady state | 2.5556632 | 0.1330766 | snoRNAs |
| SNORD10 | A tail | steady state | 2.5150986 | 0.07237 | snoRNAs |
| SNORD83A | A tail | steady state | 2.4477584 | 0.2317469 | snoRNAs |
| SNORA33 | A tail | steady state | 2.4123692 | 0.7381494 | snoRNAs |

| | | | | | |
|---|---|---|---|---|---|
| RNVU1-31 | A tail | steady state | 2.3473352 | 0.2803267 | snRNAs |
| VTRNA1-2 | A tail | steady state | 2.29308 | 1.1317593 | Pol3 RNAs |
| SNORD37 | A tail | steady state | 2.2909602 | 1.1437902 | snoRNAs |
| SNORA50B | A tail | steady state | 2.2740271 | 0.2081188 | snoRNAs |
| SNORA3A | A tail | steady state | 2.17441 | 0.1226436 | snoRNAs |
| SNORA70 | A tail | steady state | 2.1590106 | 0.2043477 | snoRNAs |
| SNORD34 | A tail | steady state | 2.09566 | 0.5238468 | snoRNAs |
| SNORD118 | A tail | steady state | 2.062694 | 0.2328296 | snRNAs |
| SNORD55 | A tail | steady state | 2.0355605 | 0.0676998 | snoRNAs |
| SNORA73A | A tail | steady state | 2.0264168 | 0.1105979 | snoRNAs |
| SNORA50C | A tail | steady state | 2.0190657 | 0.0670993 | snoRNAs |
| SCARNA11 | A tail | steady state | 2.0022332 | 0.4177113 | snoRNAs |
| SNORA74B | A tail | steady state | 1.973142 | 0.1246197 | snoRNAs |
| SCARNA7 | A tail | steady state | 1.8772894 | 1.1112327 | snoRNAs |
| SNORA52 | A tail | steady state | 1.8614033 | 0.1012285 | snoRNAs |
| SNORA68 | A tail | steady state | 1.8386847 | 0.1413209 | snoRNAs |
| SNORA10 | A tail | steady state | 1.8226246 | 0.1755484 | snoRNAs |
| SNORD95 | A tail | steady state | 1.8206765 | 0.4788533 | snoRNAs |
| SNORD13 | A tail | steady state | 1.7263186 | 0.1214848 | snoRNAs |
| SNORA69 | A tail | steady state | 1.7028986 | 1.0432015 | snoRNAs |
| RNA5S | A tail | steady state | 1.6698503 | 0.0490049 | Pol3 RNAs |
| SNORA80C | A tail | steady state | 1.6417663 | 0.2721396 | snoRNAs |
| SNORA37 | A tail | steady state | 1.5526334 | 0.2225888 | snoRNAs |
| RNVU1-7 | A tail | steady state | 1.5001389 | 0.4291496 | snRNAs |
| SNORA2C | A tail | steady state | 1.4995592 | 0.1958016 | snoRNAs |
| SCARNA4 | A tail | steady state | 1.4890652 | 0.5672282 | snoRNAs |
| SNORD67 | A tail | steady state | 1.4883403 | 0.2347288 | snoRNAs |
| SNORD16 | A tail | steady state | 1.478845 | 0.6004506 | snoRNAs |
| SNORA14B | A tail | steady state | 1.4685985 | 0.203816 | snoRNAs |
| SNORA21B | A tail | steady state | 1.4601572 | 0.495087 | snoRNAs |
| RNVU1-14 | A tail | steady state | 1.4216541 | 0.5408235 | snRNAs |
| SNORA71D | A tail | steady state | 1.4196864 | 0.1043089 | snoRNAs |
| SNORA71A | A tail | steady state | 1.3991876 | 0.1634114 | snoRNAs |
| SNORA71C | A tail | steady state | 1.3757187 | 0.0962043 | snoRNAs |
| SNORA73B | A tail | steady state | 1.2622906 | 0.1069039 | snoRNAs |
| SNORA16A | A tail | steady state | 1.22739 | 0.2154371 | snoRNAs |
| SNORD32A | A tail | steady state | 1.158819 | 0.108048 | snoRNAs |
| RNU1 WT | A tail | steady state | 1.1468348 | 0.0388648 | snRNAs |
| RNU5B-1 | A tail | steady state | 1.0636861 | 0.0794291 | snRNAs |
| SNORA71E | A tail | steady state | 1.0294547 | 0.2877386 | snoRNAs |
| RNU5A-1 | A tail | steady state | 1.0200834 | 0.0788905 | snRNAs |
| SNORA23 | A tail | steady state | 0.8622111 | 0.0989171 | snoRNAs |
| RNU5F-1 | A tail | steady state | 0.8096508 | 0.1191262 | snRNAs |

| | | | | | |
|---|---|---|---|---|---|
| RNU4ATAC | A tail | steady state | 0.7131818 | 0.3690658 | snRNAs |
| RNU2-2P | A tail | steady state | 0.6247168 | 0.4188405 | snRNAs |
| SNORD17 | A tail | steady state | 0.537848 | 9.98E-03 | snoRNAs |
| RNU4-2 | A tail | steady state | 0.4042776 | 0.0560344 | snRNAs |
| RNU5D-1 | A tail | steady state | 0.357991 | 0.1971877 | snRNAs |
| RNU4-1 | A tail | steady state | 0.2376747 | 0.036571 | snRNAs |
| RNU5E-1 | A tail | steady state | 0.1984127 | 0.1984127 | snRNAs |
| SNORA3B | A tail | steady state | 0.1818422 | 0.0491628 | snoRNAs |
| RNU12 | A tail | steady state | 0.1237855 | 3.14E-03 | snRNAs |
| AL049766.1 | A tail | steady state | 0 | 0 | snoRNAs |
| AL353804.6 | A tail | steady state | 0 | 0 | snoRNAs |
| RNA5-8S | A tail | steady state | 0 | 0 | Pol3 RNAs |
| RNA5SP515 | A tail | steady state | 0 | 0 | Pol3 RNAs |
| RNU2-63P | A tail | steady state | 0 | 0 | snRNAs |
| SNORD97 | A tail | steady state | 0 | 0 | snoRNAs |
| RNVU1-1 | U tail | steady state | 11.9191919 | 4.1005622 | snRNAs |
| RNVU1-6 | U tail | steady state | 10.512807 | 3.2635467 | snRNAs |
| RNVU1-15 | U tail | steady state | 9.6617668 | 0.9618501 | snRNAs |
| VTRNA1-3 | U tail | steady state | 7.6125118 | 0.5628641 | Pol3 RNAs |
| VTRNA1-1 | U tail | steady state | 5.4586375 | 0.8720729 | Pol3 RNAs |
| VTRNA1-2 | U tail | steady state | 4.1102994 | 0.450783 | Pol3 RNAs |
| RNY3 | U tail | steady state | 3.766729 | 0.3587182 | Pol3 RNAs |
| RNVU1-27 | U tail | steady state | 3.3706227 | 0.4189543 | snRNAs |
| RNVU1-14 | U tail | steady state | 2.7549342 | 1.8648915 | snRNAs |
| RNU4ATAC | U tail | steady state | 2.4020125 | 0.8562795 | snRNAs |
| RNA5SP452 | U tail | steady state | 2.3809524 | 2.3809524 | Pol3 RNAs |
| RNU5D-1 | U tail | steady state | 2.3437817 | 0.2233464 | snRNAs |
| SNORD68 | U tail | steady state | 2.2373394 | 0.4713669 | snoRNAs |
| RNU2 WT | U tail | steady state | 1.7446028 | 0.079809 | snRNAs |
| RNA5-8S | U tail | steady state | 1.5873016 | 1.5873016 | Pol3 RNAs |
| RNY1 | U tail | steady state | 1.5604116 | 0.2865397 | Pol3 RNAs |
| RNVU1-31 | U tail | steady state | 1.4179262 | 0.1737411 | snRNAs |
| SNORD105B | U tail | steady state | 1.3806755 | 0.3319139 | snoRNAs |
| RNA5SP304 | U tail | steady state | 1.010101 | 1.010101 | Pol3 RNAs |
| RNVU1-7 | U tail | steady state | 0.7975549 | 0.2306013 | snRNAs |
| RNU1 WT | U tail | steady state | 0.7399736 | 0.0155739 | snRNAs |
| SCARNA21B | U tail | steady state | 0.5910037 | 0.4102116 | snoRNAs |
| RNU6ATAC | U tail | steady state | 0.589198 | 0.0960173 | Pol3 RNAs |
| SNORD67 | U tail | steady state | 0.5742763 | 0.0561498 | snoRNAs |
| SNORA4 | U tail | steady state | 0.5085566 | 0.0967909 | snoRNAs |
| SNORA9 | U tail | steady state | 0.4776784 | 0.2411097 | snoRNAs |
| RNU5A-1 | U tail | steady state | 0.4400321 | 0.0824041 | snRNAs |
| SNORA71E | U tail | steady state | 0.4397537 | 0.4397537 | snoRNAs |

| | | | | | |
|---|---|---|---|---|---|
| SNORA23 | U tail | steady state | 0.4297525 | 0.0478662 | snoRNAs |
| SNORD83B | U tail | steady state | 0.4292869 | 0.0114373 | snoRNAs |
| SNORD23 | U tail | steady state | 0.4193253 | 0.0883972 | snoRNAs |
| SNORA38 | U tail | steady state | 0.4090129 | 0.2134514 | snoRNAs |
| SNORA52 | U tail | steady state | 0.4074387 | 6.43E-03 | snoRNAs |
| RNU5B-1 | U tail | steady state | 0.4040963 | 0.0519102 | snRNAs |
| SNORA16A | U tail | steady state | 0.4019028 | 0.2170372 | snoRNAs |
| SNORA71C | U tail | steady state | 0.3894795 | 0.0114525 | snoRNAs |
| SNORD95 | U tail | steady state | 0.3766954 | 0.2071215 | snoRNAs |
| SNORD16 | U tail | steady state | 0.3756704 | 0.1181019 | snoRNAs |
| SNORA71D | U tail | steady state | 0.3733102 | 0.0131134 | snoRNAs |
| SNORA77 | U tail | steady state | 0.3696459 | 0.0470144 | snoRNAs |
| SNORA38B | U tail | steady state | 0.3665896 | 0.2475177 | snoRNAs |
| SNORA50B | U tail | steady state | 0.3379951 | 0.1380685 | snoRNAs |
| SNORD37 | U tail | steady state | 0.3354467 | 0.2039743 | snoRNAs |
| RNU6 WT | U tail | steady state | 0.3354158 | 0.0744967 | Pol3 RNAs |
| SNORD10 | U tail | steady state | 0.3293936 | 8.31E-03 | snoRNAs |
| SNORA30 | U tail | steady state | 0.3236246 | 0.3236246 | snoRNAs |
| SNORA80B | U tail | steady state | 0.3236134 | 0.0657561 | snoRNAs |
| SNORA57 | U tail | steady state | 0.3132116 | 0.0641147 | snoRNAs |
| SNORA71A | U tail | steady state | 0.3084307 | 0.0136818 | snoRNAs |
| SNORA80A | U tail | steady state | 0.3064984 | 0.0617115 | snoRNAs |
| SCARNA22 | U tail | steady state | 0.3044528 | 0.0243796 | snoRNAs |
| SNORA50C | U tail | steady state | 0.2945456 | 0.0210607 | snoRNAs |
| SNORA77B | U tail | steady state | 0.2873215 | 0.0140764 | snoRNAs |
| SNORA17B | U tail | steady state | 0.2833723 | 0.0241269 | snoRNAs |
| SCARNA4 | U tail | steady state | 0.2804049 | 0.1631066 | snoRNAs |
| AL138963.2 | U tail | steady state | 0.2773572 | 0.0793381 | snoRNAs |
| SNORA73B | U tail | steady state | 0.2702016 | 0.0290998 | snoRNAs |
| SNORA51 | U tail | steady state | 0.2687125 | 0.0781667 | snoRNAs |
| SNORA47 | U tail | steady state | 0.2681621 | 0.0121519 | snoRNAs |
| RNU2-2P | U tail | steady state | 0.2614392 | 0.1974507 | snRNAs |
| RNU12 | U tail | steady state | 0.2505934 | 0.015197 | snRNAs |
| SNORA3B | U tail | steady state | 0.2448682 | 0.0503994 | snoRNAs |
| SCARNA12 | U tail | steady state | 0.2427116 | 0.015322 | snoRNAs |
| SNORD88A | U tail | steady state | 0.2344371 | 0.0284634 | snoRNAs |
| SNORD88C | U tail | steady state | 0.2337579 | 0.02119 | snoRNAs |
| SNORD97 | U tail | steady state | 0.2314815 | 0.2314815 | snoRNAs |
| SNORA68 | U tail | steady state | 0.2289994 | 0.0109145 | snoRNAs |
| SNORD94 | U tail | steady state | 0.2236243 | 0.0630922 | snoRNAs |
| SNORA5C | U tail | steady state | 0.2222237 | 0.0459954 | snoRNAs |
| SNORA78 | U tail | steady state | 0.2154119 | 1.53E-03 | snoRNAs |
| SNORA48 | U tail | steady state | 0.2122808 | 0.0178264 | snoRNAs |

| | | | | | |
|---|---|---|---|---|---|
| SNORD17 | U tail | steady state | 0.2091592 | 0.015888 | snoRNAs |
| RNU4-1 | U tail | steady state | 0.2066143 | 0.0170087 | snRNAs |
| SNORD83A | U tail | steady state | 0.2014602 | 0.1028912 | snoRNAs |
| TERC | U tail | steady state | 0.1971595 | 0.0163126 | snRNAs |
| SNORD118 | U tail | steady state | 0.19334 | 0.1014049 | snRNAs |
| SNORA74B | U tail | steady state | 0.192123 | 0.192123 | snoRNAs |
| SCARNA2 | U tail | steady state | 0.1904762 | 0.1904762 | snRNAs |
| RNU5F-1 | U tail | steady state | 0.1878715 | 0.0443952 | snRNAs |
| SNORD104 | U tail | steady state | 0.1858144 | 0.1026173 | snoRNAs |
| SNORD33 | U tail | steady state | 0.179208 | 0.052983 | snoRNAs |
| SNORA64 | U tail | steady state | 0.1761341 | 0.0180817 | snoRNAs |
| SCARNA20 | U tail | steady state | 0.163582 | 0.0327445 | snoRNAs |
| SNORA5A | U tail | steady state | 0.1633987 | 0.1633987 | snoRNAs |
| SNORA80C | U tail | steady state | 0.1622743 | 0.0316719 | snoRNAs |
| SNORD55 | U tail | steady state | 0.1609567 | 0.0480512 | snoRNAs |
| SCARNA10 | U tail | steady state | 0.1602564 | 0.1602564 | snoRNAs |
| SNORA21B | U tail | steady state | 0.1565898 | 0.1043221 | snoRNAs |
| RN7SK | U tail | steady state | 0.1509768 | 0.0190392 | Pol3 RNAs |
| RNU4-2 | U tail | steady state | 0.1488963 | 0.025386 | snRNAs |
| SNORA14B | U tail | steady state | 0.1415428 | 0.1415428 | snoRNAs |
| SNORD22 | U tail | steady state | 0.1406164 | 0.0341482 | snoRNAs |
| SNORA70 | U tail | steady state | 0.1388354 | 0.0364079 | snoRNAs |
| SNORA61 | U tail | steady state | 0.1365914 | 0.0642052 | snoRNAs |
| SNORA37 | U tail | steady state | 0.130719 | 0.130719 | snoRNAs |
| SNORA18 | U tail | steady state | 0.129199 | 0.129199 | snoRNAs |
| SNORA67 | U tail | steady state | 0.1281864 | 0.0912911 | snoRNAs |
| SNORA80D | U tail | steady state | 0.1196026 | 0.025092 | snoRNAs |
| RPPH1 | U tail | steady state | 0.1168225 | 0.0127829 | Pol3 RNAs |
| SNORA21 | U tail | steady state | 0.1152982 | 0.0576549 | snoRNAs |
| RNU11 | U tail | steady state | 0.1104997 | 0.0255615 | snRNAs |
| SNORA74D | U tail | steady state | 0.1007049 | 0.1007049 | snoRNAs |
| SNORA80E | U tail | steady state | 0.0969552 | 0.0491603 | snoRNAs |
| SNORD13 | U tail | steady state | 0.0957736 | 5.04E-03 | snoRNAs |
| SNORA2C | U tail | steady state | 0.094162 | 0.094162 | snoRNAs |
| SNORA65 | U tail | steady state | 0.0879507 | 0.0513994 | snoRNAs |
| SNORA7B | U tail | steady state | 0.0821076 | 0.0296715 | snoRNAs |
| SNORA10 | U tail | steady state | 0.0801976 | 0.0114281 | snoRNAs |
| SNORA63 | U tail | steady state | 0.0766284 | 0.0766284 | snoRNAs |
| RMRP | U tail | steady state | 0.0753086 | 2.36E-03 | Pol3 RNAs |
| SCARNA8 | U tail | steady state | 0.0739905 | 0.0454583 | snoRNAs |
| SNORD46 | U tail | steady state | 0.0726216 | 0.0726216 | snoRNAs |
| SNORD86 | U tail | steady state | 0.0706215 | 0.0706215 | snoRNAs |
| SNORD3 | U tail | steady state | 0.062456 | 0.0232178 | snRNAs |

| | | | | | |
|---|---|---|---|---|---|
| SNORA7A | U tail | steady state | 0.0618356 | 6.83E-03 | snoRNAs |
| SNORA73A | U tail | steady state | 0.0582863 | 9.13E-03 | snoRNAs |
| SNORD32A | U tail | steady state | 0.0543596 | 0.0543596 | snoRNAs |
| SCARNA11 | U tail | steady state | 0.050543 | 0.0284698 | snoRNAs |
| RNA5S | U tail | steady state | 0.0403309 | 3.11E-03 | Pol3 RNAs |
| RN7SL1 | U tail | steady state | 0.0354999 | 2.58E-03 | Pol3 RNAs |
| RN7SL2 | U tail | steady state | 0.0347327 | 3.11E-03 | Pol3 RNAs |
| SNORA3A | U tail | steady state | 0.027674 | 0.027674 | snoRNAs |
| RN7SL3 | U tail | steady state | 0.0260796 | 3.73E-03 | Pol3 RNAs |
| SNORA84 | U tail | steady state | 0.0211154 | 0.0124392 | snoRNAs |
| SNORA44 | U tail | steady state | 0.0142572 | 0.0142572 | snoRNAs |
| AC090227.1 | U tail | steady state | 0 | 0 | snoRNAs |
| AL049766.1 | U tail | steady state | 0 | 0 | snoRNAs |
| AL353804.6 | U tail | steady state | 0 | 0 | snoRNAs |
| RNA5SP19 | U tail | steady state | 0 | 0 | Pol3 RNAs |
| RNA5SP515 | U tail | steady state | 0 | 0 | Pol3 RNAs |
| RNU2-63P | U tail | steady state | 0 | 0 | snRNAs |
| RNU5E-1 | U tail | steady state | 0 | 0 | snRNAs |
| SCARNA5 | U tail | steady state | 0 | 0 | snoRNAs |
| SCARNA7 | U tail | steady state | 0 | 0 | snoRNAs |
| SNORA33 | U tail | steady state | 0 | 0 | snoRNAs |
| SNORA62 | U tail | steady state | 0 | 0 | snoRNAs |
| SNORA69 | U tail | steady state | 0 | 0 | snoRNAs |
| SNORD24 | U tail | steady state | 0 | 0 | snoRNAs |
| SNORD34 | U tail | steady state | 0 | 0 | snoRNAs |
| SNORD36B | U tail | steady state | 0 | 0 | snoRNAs |
| SNORD38A | U tail | steady state | 0 | 0 | snoRNAs |
| SNORD66 | U tail | steady state | 0 | 0 | snoRNAs |

**Supplementary Table S6.** Mean terminal 3' uridine lengths of steady state and nascent Pol-III RNAs.

| genename | steady state | nascent | delta | pvalue |
|---|---|---|---|---|
| RNU6 WT | 4.215914 | 3.566581 | 0.649333 | 2.586449 |
| RNU6ATAC | 3.743753 | 3.327425 | 0.416328 | 0.946156 |
| RN7SK | 2.527791 | 2.540313 | -0.01252 | 0.138962 |
| RN7SL2 | 1.340933 | 1.493261 | -0.15233 | 2.196118 |
| RNA5S | 2.209023 | 2.394953 | -0.18593 | 1.912702 |
| RN7SL3 | 1.800964 | 1.993557 | -0.19259 | 1.503711 |
| RN7SL1 | 1.508686 | 1.70268 | -0.19399 | 2.268435 |
| RNY1 | 2.144575 | 2.82461 | -0.68004 | 1.286159 |
| VTRNA1-2 | 3.352802 | 4.043417 | -0.69062 | 0.729326 |
| RPPH1 | 2.066677 | 2.963807 | -0.89713 | 1.431756 |
| RNY3 | 2.42774 | 3.465527 | -1.03779 | 2.10345 |
| VTRNA1-1 | 3.479397 | 4.667513 | -1.18812 | 1.199403 |
| RMRP | 1.173807 | 3.150638 | -1.97683 | 2.808501 |
| VTRNA1-3 | 6.636212 | 9.457093 | -2.82088 | 0.933025 |
